# SURVIVIN IN SYNERGY WITH BAF/SWI COMPLEX BINDS BIVALENT CHROMATIN REGIONS AND ACTIVATES DNA DAMAGE RESPONSE IN CD4+ T CELLS

**DOI:** 10.1101/2024.03.05.583464

**Authors:** Venkataragavan Chandrasekaran, Karin M.E. Andersson, Malin Erlandsson, Shuxiang Li, Maria-Jose Garcia-Bonete, Eric Malmhäll-Bah, Johansson Pegah, Gergely Katona, Maria I. Bokarewa

## Abstract

This study explores a regulatory role of oncoprotein survivin on the bivalent regions of chromatin (BvCR) characterized by concomitant deposition of trimethylated lysine of histone H3 at position 4 (H3K4me3) and 27 (H3K27me3).

Intersect between BvCR and chromatin sequences bound to survivin demonstrated their co-localization on *cis*-regulatory elements of genes which execute DNA damage control in primary human CD4^+^ cells. Survivin anchored BRG1-complex to BvCR to repress DNA damage repair genes in IFNγ-stimulated CD4^+^ cells. In contrast, survivin inhibition shifted the functional balance of BvCR in favor of H3K4me3, which activated DNA damage recognition and repair. Co-expression of BRG1, survivin and IFNγ in CD4^+^ cells of patients with rheumatoid arthritis identified arthritogenic BRG1^hi^ cells abundant in autoimmune synovia. Immunomodulating drugs inhibited the subunits anchoring BRG1-complex to BvCR, which changed the arthritogenic profile.

Together, this study demonstrates the function of BvCR in DNA damage control of CD4^+^ cells offering an epigenetic platform for survivin and BRG1-complex targeting interventions to combat autoimmunity.

**Summary:** This study shows that bivalent chromatin regions accommodate survivin which represses DNA repair enzymes in IFNγ-stimulated CD4^+^ T cells. Survivin anchors BAF/SWI complex to these regions and supports autoimmune profile of T cells, providing novel targets for therapeutic intervention.

## Introduction

The octamer of histone proteins and DNA strands form the nucleosome, the basic repeated unit of chromatin. The N-terminal tails of histone proteins protrude from the nucleosome and regulate chromatin function (Janssen and Lorincz, 2021; Taylor and Young, 2021). The combination of histone H3 tail modifications trimethylated lysine 4 (H3K4me3) and lysine 27 (H3K27me3) define bivalent chromatin regions (BvCR) across the genome (Bernstein et al., 2006; Blanco et al., 2020).

BvCR are well-studied in developmental biology wherein, spatiotemporal control of developmental stages of a cell is achieved by resolution of the BvCR into active, dominated by H3K4me3, or repressive, dominated by H3K27me3 regions (Bernstein et al., 2006). In terminally differentiated CD4^+^ cells of the peripheral blood, resolution of bivalency is not always observed, which maintains flexibility of gene expression in response to the surrounding immune environment (Wei et al., 2009). Mathematical modelling (Sneppen and Ringrose, 2019; Zhao et al., 2021) and experimental evidence (Kinkley et al., 2016) indicate duality of BvCR where both H3K4me3 and H3K27me3 can stably exist on one or both tails of a single nucleosome to flexibly finetune gene transcription. Deposition of BvCR is mediated by catalytically active subunits of chromatin remodeling complexes including BRG1 associated SWItch/Sucrose Non-Fermentable complex (BAF/SWI), Polycomb Repressive Complex 2 (PRC2), and Complex of Proteins Associated with Set1 (COMPASS). BvCR have been described in several immunocompetent cells including CD4^+^ T cells (Harker et al., 2011; Kinkley et al., 2016; Roh et al., 2006; Wei et al., 2009), B cells (Bossen et al., 2015), adipocytes (Park et al., 2021), and fibroblasts (Alver et al., 2017). However, the precise function of BvCR in terminally differentiated cells is still debated.

In addition to transcriptional control, histone modifications are critical to the maintenance of genome integrity within the nucleus. Involvement of H3K4me3 in DNA damage recognition has been proposed based on its enrichment in chromosomal break sites (Faucher and Wellinger, 2010; Barski et al., 2007). Different DNA repair mechanisms are activated in response to type of DNA damage. These include the double-strand break repair by means of homologous recombination and non-homologous end joining, inter-strand cross-link repair, and the single-strand break repair by base and nucleotide excision repair, mismatch repair, and trans-lesion synthesis. Natural progression of the DNA repair requires the concerted action of histone-modifying enzymes, and the chromatin remodeling complexes (Harrod et al., 2020). Deposition of histone variant H2AX phosphorylated at serine residue 139 (γH2AX) on DNA is routinely used to identify the sites of DNA double-strand breaks. γH2AX acts as a platform for recruitment of DNA repair complexes of the MRE11-RAD50-NBS1 (MRN) proteins and NuA4/TIP60 acetyltransferase. Abnormal DNA repair can impact cell cycle regulation. For example, deficiency in the repair protein MRE11 in naïve and memory CD4^+^ cells from RA patients displayed upregulated cyclin-dependent kinase inhibitors p16, p21, and downregulated p53, characteristic for pro-inflammatory senescent phenotype (Li et al., 2016). Similarly, deficiency of the ATM (ataxia telangiectasia mutated) enzyme has been tightly linked to the impaired DNA repair in RA CD4^+^ cells (Shao, 2018; Lee and Paull, 2021). While histone methylation abundance contributing to DNA repair have been documented (Pai et al., 2014; Jeon et al., 2020), a role of BvCR in DNA repair has not been previously reported.

Survivin is an inhibitor of apoptosis protein which is best known for its role in cell cycle control through its interaction with phosphorylated threonine-3 of histone H3 (Yamagishi et al., 2010; Niedzialkowska et al., 2012) as part of the Chromosomal Passenger Complex (Jeyaprakash et al., 2007; Carmena et al., 2012). It is encoded by the gene *BIRC5*. Cytoplasmic localization of survivin leads to anti-apoptotic activity (Altieri, 2003), while nuclear survivin acts in gene transcription through association with transcription factors (TF, Fukuda et al., 2015; Wang et al., 2010; Erlandsson et al., 2022). Survivin is important for the maturation of lymphocytes (Kornacker et al., 2001; Okada et al., 2004; Song et al., 2005; Xing et al., 2003; Andersson et al., 2015), and hematopoietic stem cells (Fukuda et al., 2002, 2015). Survivin deletion in early thymocytes prevents formation of functional T cell receptor and obstructs T cell development (Okada et al., 2004; Xing et al., 2003). In mature CD4^+^ cells, survivin is essential for maintaining the effector activity of T cells contributing to IFNγ signaling which is critically involved in autoimmune inflammation (Song et al., 2005; Erlandsson et al., 2022). Survivin and cell death inhibition have been connected by anti-apoptotic properties of survivin interaction with caspases in cytoplasm and Smac/Diablo in mitochondria (Gravina et al., 2017).

Interaction of survivin with threonine 3 of histone H3 (Yamagishi et al., 2010; Niedzialkowska et al., 2012), together with our recent findings that survivin interacts with the catalytic subunit of the PRC2 repressive complex and inhibits trimethylation of histone H3K27 (Jensen et al., 2023), prompted us to investigate if survivin bound to the BvCR in CD4^+^ cells and if this binding effected CD4^+^ cell function. To examine this, we performed parallel chromatin sequencing in primary human CD4^+^ cells to annotate survivin to the BvCR and to explore the TF landscape of those BvCR to identify chromatin modifying complexes. Molecular docking approach and composition-based machine learning of suitable survivin binding regions was used to model survivin and BAF/SWI complex interaction in proximity of nucleosome. Further, functional studies on survivin-dependent histone H3 modifications and the transcriptome of CD4^+^ cells suggest BvCR involvement to the DNA damage response mediated by the BAF/SWI complex. Finally, we investigated the role of survivin and BAF/SWI complex in autoimmune CD4^+^ cells of patients with rheumatoid arthritis, and the ability of anti-rheumatic drugs to modulate the BAF/SWI complex composition.

## Results

### 1. Survivin colocalizes with histone H3 marks in BvCR

Sequencing of chromatin regions containing deposition of H3K4me3, H3K27me3, and H3K27ac in CD4^+^ cells, revealed an overlap of non-redundant peaks in a total of 6199 regions across the genome, further called bivalent chromatin regions (BvCR) (**Figure 1A**). In identical experimental settings, we identified genome location of survivin peaks (n=13703, **Figure 1A**) by sequencing chromatin immunoprecipitated with survivin in four independent experiments. Integrating BvCR with survivin-bound chromatin regions (**Figure 1B**), we observed that 65% of BvCR contained survivin peaks (survivin-positive BvCR, S+BvCR, 4068/6199 regions) (**Figure 1C**). The association between survivin and BvCR is not mutual since a large fraction of survivin peaks (n=7823, 57%) are also localized apart from BvCR. We found that, H3K4me3 mark dominated the BvCR (43%, H3K4me3-BvCR), followed by H3K27me3 (33%, H3K27me3-BvCR) and H3K27ac (24%, H3K27ac-BvCR) (**Figure 1C**, **Supplementary Figure S1A**). This frequency distribution of BvCR by the dominant H3 mark was comparable for the BvCR and S+BvCR (**Figure 1C**).

**Figure 1.**
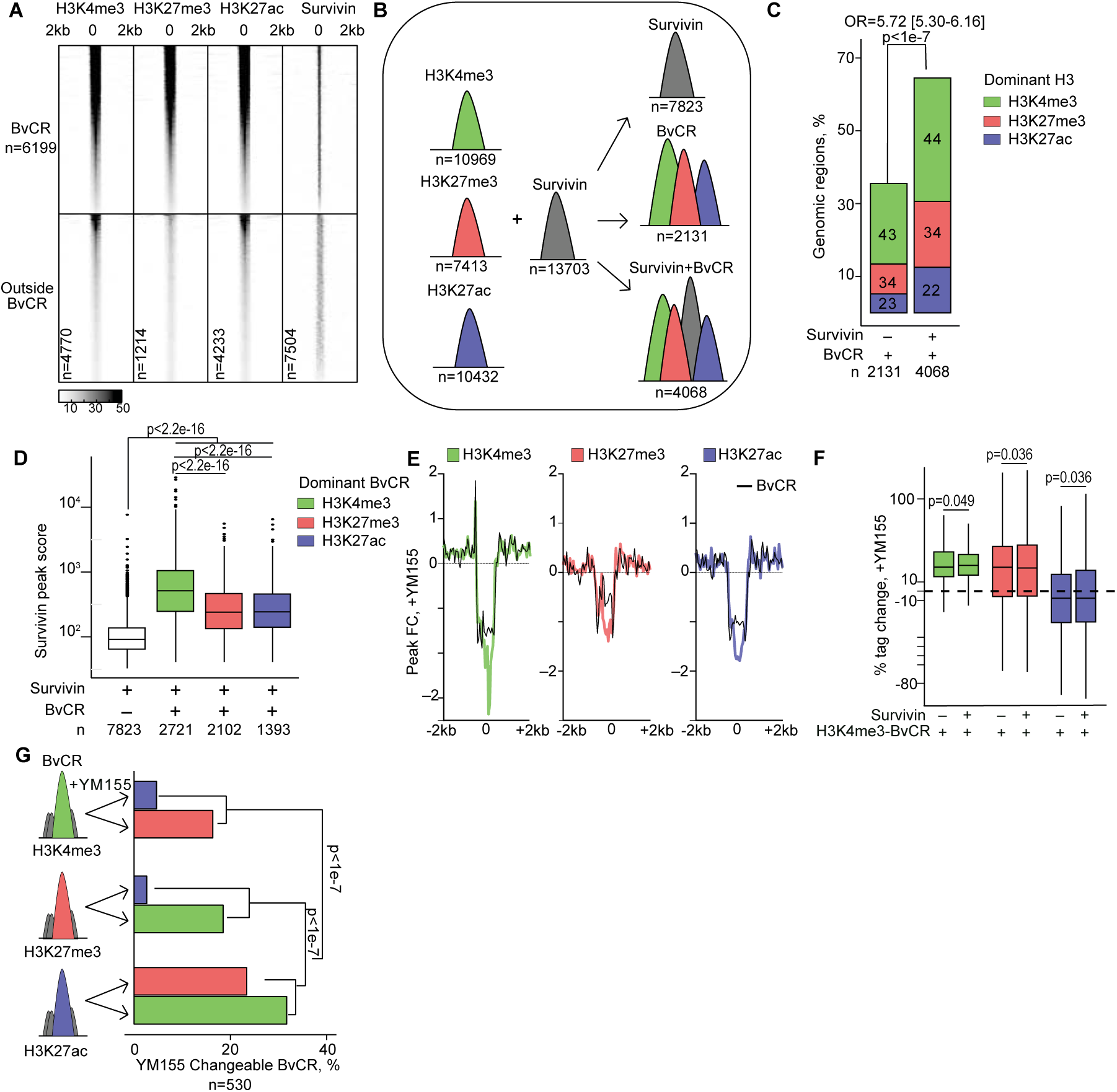
Survivin accumulates in H3K4me3-dominant bivalent chromatin regions in primary CD4 cells. (A) Heatmap of ChIP-seq peaks within and outside the bivalent chromatin regions (BvCR) defined by genomic overlap between the histone H3-marks. (B) Cartoon of BvCR and survivin deposition. (C) Frequency difference of BvCR without and with survivin. Chi-square test p-value is shown. Numbers within bars indicate percentage of BvCR dominant in individual H3 marks. (D) Box plots of survivin peak scores within BvCR dominant by H3K4me3, H3K27me3 and H3K27ac marks. Kolmogorov-Smirnov test p-values are shown. (E) Histogram of fold change in mean H3 peak score in naïve and YM155-treated CD4+ cells within (black) and outside BvCR (colored). (F) Box plot of H3 tag change within H3K4me3 dominant BvCR, after YM155 treatment. Mann-Whitney test p-values are indicated. (G) Frequency of changeable BvCR that shifts in dominant H3 after YM155 treatment. Kolmogorov-Smirnov test p-values are shown.

Next, we investigated the strength of the survivin peaks within BvCR and found that the average survivin peak score was higher if the peaks co-localized with BvCR, compared to genomic regions that contained survivin peaks only. Besides, deposition of survivin within BvCR was significantly larger in the H3K4me3-BvCR (**Figure 1D**). Similarly, score of the H3 peaks showed significant differences between those localized within S+BvCR and the H3 peaks not confined to survivin or BvCR colocalization (**Supplementary Figure S1B**), with H3K4me3-BvCR having the highest peak scores. Together, these results demonstrated non-randomness of survivin binding to chromatin by frequently annotating to BvCR across the genome, where deposition of survivin was associated with a reciprocal increase in H3 peaks in BvCR, appreciably within the H3K4me3-dominant BvCR.

### 2. Survivin binds H3K4me3 dominant BvCR and regulates its functional status

Searching for understanding of survivin function within BvCR, we asked if the deposition of individual histone H3 marks was influenced by survivin. To investigate this, we cultured CD4^+^T cells in presence of the survivin inhibitor YM155 and performed the chromatin sequencing analysis of H3K4me3, H3K27me3 and H3K27ac deposition. The adjusted average enrichment profile of BvCR showed that YM155-treated cells altered the deposition of all three H3 marks. The difference was clearly seen in the peak center of individual H3 marks (**Figure 1E**). Further, we analyzed if deposition of survivin within the BvCR affected response to YM155 treatment. We observed that the presence of survivin within BvCR led to a YM155-mediated increase in H3 mark deposition only within H3K4me3-BvCR (**Figure 1F**), and only within the H3K4me3 mark in the other dominant BvCR (**Supplementary Figure S1C**) where H3K4me3 had lower peak score prior to YM155 treatment. The quantitative increase in H3 tag deposition caused a shift in the dominant H3 mark within BvCR, which we termed as changeable BvCR (**Figure 1G**). In total, such a shift occurred in 530 BvCR (8.55%) and was less prevalent among the S+BvCR (325/4068 vs 205/2131, p-value=0.03). We observed that the BvCR dominant in either H3K4me3 or H3K27me3 shifted into each other in equal frequency, reflecting the functional bivalency of those chromatin regions (**Figure 1G**). Additional support to the bivalency was provided by the H3K27ac-BvCR, which frequently lost their status and gained the dominance of H3K4me3 (32%), followed by H3K27me3 (23%). Therefore, the analysis of YM155-treated CD4^+^ cells showed that survivin inhibition increased the density of the lysine trimethylation on histone H3, largely increasing the proportion of H3K4me3-BvCR.

### 3. BvCR dominated by H3K4me3 control the DNA damage response through BAF/SWI complex

To connect H3 tag deposition in BvCR and long-distance gene regulation, we exploited the experimentally confirmed connections between *cis*-RE and genes publicly available through the GeneHancer database (Fishilevich et al., 2017). In total, we observed that between 59-65% of BvCR were located within *cis*-RE (**Figure 2A**). Further focusing on the transcriptome of CD4^+^ cells, we identified 4212 protein-coding genes connected to those *cis*-RE and whose expression could be regulated by the BvCR located within *cis*-RE (**Figure 2A**). The activating H3K4me3-BvCR had the largest number of connected genes, reflecting the fact that we analyzed genes actively transcribed within CD4^+^ cells (**Figure 2A**). We asked if functional changes in the BvCR translated into changes in transcription of the connected genes, by investigating which of the connected genes were sensitive to IFNγ stimulation or survivin inhibition by YM155.

**Figure 2.**
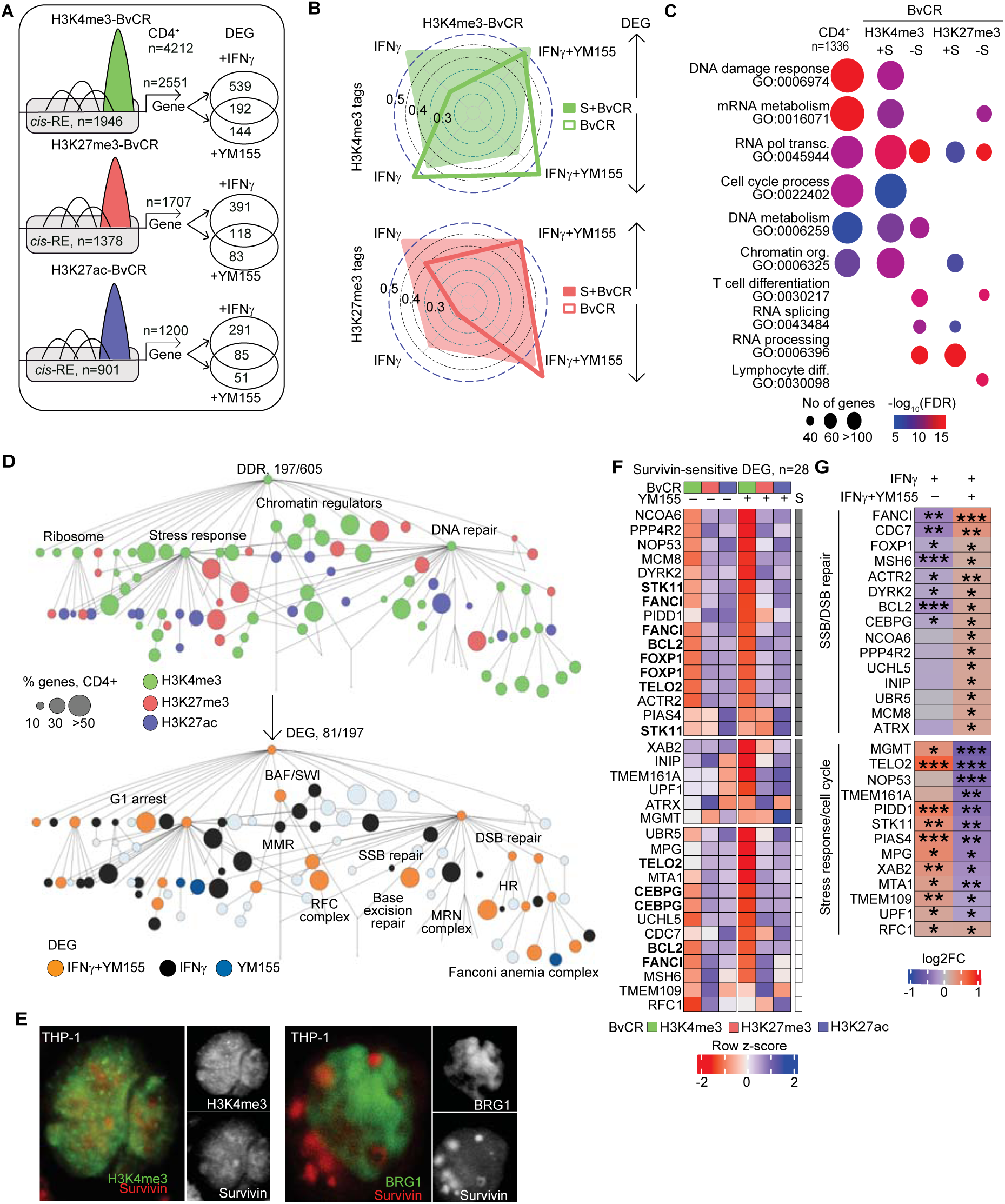
BvCR dominant in H3K4me3 together with survivin regulate transcription of DNA damage response genes. (A) Cartoon of analysis strategy. BvCR within genomic regulatory elements (*cis*-RE, grey boxes) connected to genes, filtered on the protein-coding genes expressed in CD4^+^ cells, by RNA-seq. Transcription difference in CD4^+^ cells treated with IFNg or IFNg+YM155 compared to sham cultures was calculated by DESeq2. Differentially expressed genes (DEG) were defined by a nominal p-value<0.05. (B) Radar plot of Spearman’s rho correlations between H3K4me3 and H3K27me3 tag deposition change in all and survivin-positive BvCR and transcription change in CD4^+^ cells treated with IFNg or IFNg+YM155. Arrows indicate direction of transcription change. (C) Bubble plot of enrichment in biological processes among CD4^+^ expressed genes connected to all and survivin-positive BvCR. Bubble size indicates protein number in the process. Color intensity shows false discovery range (FDR). (D) Nodes of the DDR network are colored by dominant H3 mark in BvCR connected to genes within nodes (top map) and by transcription change after IFNγ or YM155 treatment (bottom map). Size of bubble corresponds to percentage of BvCR-connected genes within each node. DDR, DNA damage response. MMR, mismatch repair. RFC, replicator factor C. SSB, single strand break. DSB, double-strand break. HR, homologous recombination. MRN, MRE11-RAD50-NBS1. (E) Confocal image of THP1 nucleus showing presence of survivin (red), nucleus (blue) and BRG1 or H3K4me3 (green). (F) Heatmap of normalized tag deposition of H3 marks, by ChIP-seq, in BvCR connected to DEG treated with IFNγ+YM155. Shaded squares indicate survivin-positive BvCR. Genes connected to multiple BvCR are marked in bold. (G) Heatmap of RNA-seq transcription difference in genes annotated to DNA repair and stress response categories. Transcription difference was calculated by DESeq2 statistics, p-values * < 0.05, ** < 0.01, *** <0.001.

Analyzing the transcriptome in CD4^+^ cells, we found that about 40% of the genes connected to any BvCR and expressed in CD4^+^ cells were differentially expressed after IFNγ and/or YM155 treatment, i.e., were IFNγ- and/or survivin-sensitive (**Figure 2A)**. Overall, changeable BvCR were less frequent in S+BvCR, while those connected to both the IFNγ-sensitive and survivin-sensitive genes were significantly predominant among S+BvCR (**Supplementary Figure S2A**) reiterating that survivin’s genomic colocalization mediated IFNγ-dependent transcription.

To investigate an internal relation between the YM155-induced changes in H3 tag deposition in BvCR and transcriptional effect on the connected genes, we built a linear regression model between these parameters in the IFNγ- and YM155-treated CD4^+^ cells compared to mock. Upregulation of IFNγ-sensitive and survivin-sensitive genes connected to survivin-positive H3K4me3-BvCR had a strong positive correlation (Spearman ρ >0.5) to the change in both H3K4me3 and H3K27me3 deposition (**Figure 2B, Supplementary Figure S2B**), suggesting that survivin deposition contributes to the dynamics of H3 tail deposition. Within H3K27me3-BvCR, the correlation between transcription and the survivin-dependent H3 tag deposition change was weaker (**Supplementary Figure S2C**). Interestingly, the correlation between a change in H3K4me3 tags (to a lesser extent in H3K27me3 tags) and downregulation of IFNγ- and survivin-sensitive genes was somewhat stronger when survivin is not present in the BvCR (**Supplementary Figure S2C**, **Figure 2B**). These findings clearly demonstrated that 1) H3K4me3 was the primary H3 modification transducing the effect of survivin accumulation in *cis-*RE into IFNγ-sensitive and survivin-sensitive upregulation of transcription, and 2) transcription of survivin-sensitive genes was dependent on a complex interplay between H3K4me3 and H3K27me3 deposition in H3K4me3-BvCR.

To better understand the cellular functions regulated by the BvCR, we searched for biological processes represented by 4212 genes connected to BvCR in CD4^+^ cells. We discovered that the DNA damage response (DDR, GO:0006974) was the principal pathway regulated by the BvCR (**Figure 2C, Supplementary Table T1**). Other biological processes enriched in the BvCR-regulated transcriptome comprised the nucleosome-modifying processes including cell cycle process (GO:0022402), mRNA metabolism (GO:0016071), RNA polymerase-dependent transcription (GO:0045944), DNA metabolism (GO:0006259), and chromatin organization (GO:0006325) (**Figure 2C**). Consistent with a strong correlation between transcription and H3K4me3 deposition change, the genes active in these six processes were predominantly connected to survivin-positive H3K4me3-BvCR. Such an enrichment was neither found in the H3K4me3-BvCR lacking survivin deposition (**Figure 2C**) nor in the H3K27me3-BvCR. On the contrary, the genes connected to H3K27me3-BvCR represented immunologically relevant processes of T cell differentiation (GO:0030217), RNA splicing (GO:0043484), RNA processing (GO:0006396) (**Figure 2C**). Further, the majority (65%) of the genes connected to H3K4me3-BvCR were annotated to any of the six enriched pathways above (**Supplementary Figure S3A**), followed by an additional subset of 30%, which was annotated to more than two of the pathways. These results suggested that H3K4me3-BvCR particularly distinguished nucleosome-modifying processes in CD4^+^T cells and primarily regulated genes of the DDR pathway.

### 4. H3K4me3-BvCR regulate the functional DDR network

To narrow down on specific functions within the DDR regulated by the H3K4me3-BvCR, we utilized the recently proposed DDR interaction network (Kratz et al., 2023) which employed a systems biology approach to catalogue protein-protein interactions and assign them into functional DDR assemblies (**Figure 2D**), comprising 605 genes in total assigned to 109 nodes. Annotation of the genes connected to the BvCR and expressed in CD4^+^ cells revealed that 197 of the genes belonged to the DDR network which were distributed between 89% of the nodes (97/109) in the DDR network (**Figure 2D, top, Supplementary Table T2**), which shows distribution of the BvCR connected genes within DDR network. To identify which H3 marks controlled specific processes within the DDR network, we calculated the mean H3 tag deposition density in the BvCR connected to the genes of each node and detected the dominant H3 mark. Among the DDR network nodes containing the BvCR connected genes, 63% were dominated by H3K4me3 tags, additional 28% had H3K27me3 tag dominance. Importantly, the core nodes of the DDR network including DNA repair, chromatin regulators, stress response and ribosome were all controlled by the H3K4me3-BvCR. The H3K4me3-BvCR nodes that contained more than half of the genes connected to the BvCR were the chromatin regulation by the BRG1-associated factor (BAF/SWI) complex (**Figure 2D, top**) that comprised the genes of *SMARCA4, SMARCE1*, *SMARCB1*, and the Replicator factor C complex comprising the genes of *RFC1, MSH2*, and *MSH6*. Gene Ontology enrichment analysis showed that more than 60% of H3K4me3-BvCR connected genes in the DDR were additionally functional in other pathways (**Supplementary Figure S3A**). Independent approaches through Gene Ontology and DDR network analysis revealed that the BAF complex subunits SMARCA4/BRG1, SMARCE1, SMARCC1, SMARCD2, SMARCB1, DPF2, and ARID1B were represented in multiple (>4) pathways, which implied that BAF/SWI complex was central for DDR and the nucleosome-modifying processes supervised by H3K4me3-BvCR. The interaction between survivin, H3K4me3-BvCR, and the BAF/SWI complex catalytic subunit BRG1 were also visualized by confocal imaging in THP1 cells known for abundant survivin expression (**Figure 2E**). Further, we noticed that processes within the categories of stress response, single-strand break (SSB) and double-strand break (DSB) repair in the DDR network were organized through H3K4me3-BvCR. The nodes controlled by H3K27me3-BvCR included homologous recombination through the MRE11-RAD50-NBS1 (MRN) complex and p53. Processes within the ribosome and stress response were distributed among all three H3 modifications and contained only a minor fraction of the genes connected to BvCR (**Figure 2D, top, Supplementary Table T2**).

Since we had demonstrated that deposition in H3K4me3 acted as a predictor of transcription change after IFNγ or YM155 treatment (**Figure 2B**), we investigated the DDR network for survivin-sensitive and IFNγ-sensitive genes. Remarkably, we found that the nodes controlled by H3K4me3 overlapped to a high extent with those containing the IFNγ- and/or survivin-sensitive genes (**Figure 2D, bottom**). Interestingly, the genes sensitive to both survivin and IFNγ were represented widely among the nodes controlled by H3K4me3-BvCR within the DDR network (**Figure 2D, bottom**), while the genes sensitive to only IFNγ were distributed among both H3K4me3-BvCR and H3K27me3-BvCR controlled nodes. Affected by IFNγ stimulation or survivin inhibition were the nodes of SSB/DSB repair including specific branches of mismatch repair (5/10 genes, *ATAD5, RFC1, CHTF18, MSH2, MSH6*), DNA replication (8/22 genes, *ATAD5, RFC1, FANCI, CHTF18, MSH2, MSH6, CTPS1, MCM8*), base excision repair (1/3 genes, *MSH2*), and nucleotide excision repair (2/4 genes, *USP7, XAB2*); Fanconi anemia complex (4/8 genes, *FANCI, RMI2, FANCA, MCM8*), and homologous recombination (4/8 genes, *MRE11, XRCC3, RMI2, UIMC1*); the G1 cell cycle arrest category (2/2 genes, *STK11, CAB39*); and stress response (44/109 genes, *USP7, ANKFY1, AMD1, RPL5, MPRIP, HSPD1, CDKN1A, PFDN5, KEAP1, CCT8, CCT8, LSS, NELFA, NELFA, RFC1, FOXK1, FOXK1, FOXK1, CUL4B, CUL4B, RIC8A, RIC8A, DPF2, DPF2, CCT2, CUL4A, XRCC3, XRCC3, CHTF18, MPG, PSMD3, MYBBP1A, MYBBP1A, TP53, CDC37, SUPT5H, LENG1, MSH2, CCT4, LARP7, DDX60, NNT, DST, DST, DST, PARP3, MSH6, PFKP, SMARCA2, AHCTF1, AHCTF1, CTPS1, PGM1, ADAR, MCM8, DCAF13, DCAF13*).

Taken together, the analysis on the biological processes controlled by BvCR in CD4^+^ cells reveal the dominant role of H3K4me3-BvCR in the core nodes of the DDR network, which spread deep into the accessory branches of the DNA repair, cell cycle, and stress response processes and potentially facilitated by the BAF/SWI complex.

### 5. Survivin inhibition counteracts IFNγ effects and triggers DNA damage recognition and repair

To underpin molecular mechanisms behind the H3K4me3-BvCR mediated regulation of DDR, we retrieved the IFNγ and/or survivin-sensitive genes connected to H3K4me3-BvCR and annotated to the DDR pathway (**Supplementary Figure S3A**). Analyzing normalized deposition of the H3 tags in the BvCR connected to the survivin-sensitive genes (n=28, **Figure 2F**), and to the IFNγ-sensitive genes (n=45, **Supplementary Figure S3B**), we found survivin deposition in the majority (63-68%) of the IFNγ- and survivin-sensitive genes connected to the BvCR. Furthermore, several of the survivin-sensitive genes (*FANCI*, *STK11*, *BCL2*, *FOXP1*, *CEBPG*) and the IFNγ-sensitive genes (*VRK1, RTEL1, DOT1L, MYC, BACH1, MDM4, AXIN2, XRCC3, HIPK2, HMGN1, BRD4, PYHIN1*) were connected to more than one H3K4me3-dominant BvCR, which multiplied survivin control. Analyzing *cis*-RE of these genes, we demonstrated that survivin inhibition with YM155 caused a change in H3K4me3 tag deposition in the corresponding BvCR (**Supplementary Figure S4**). The dominance of H3K27me3 or H3K27ac in BvCR connected to the DDR genes was infrequent, but it comprised control over *cis-*RE connected to *MRE11*, *MSH2*, *FANCA* genes essential for DNA repair. Changeable H3K4me3-BvCR were observed in 32% of the survivin-sensitive genes (9/28, *TMEM109, STK11, XAB2, RFC1, UPF1, MGMT, ATRX, INIP, TMEM161A*) (**Figure 2F**) and somewhat less frequent in the IFNγ-sensitive genes (18%, 8/45, *FANCA, RMI2, CD44, DOT1L, EXO5, SMARCAL1, MYC, CUL4A*) (**Supplementary Figure S3B**).

In contrast to the whole BvCR-connected transcriptome, a minor part of changeable H3K4me3-BvCR were connected to the DDR, indicating a stability of the DDR-related H3K4me3 mark after survivin inhibition. Quantifying the normalized tag deposition within the H3K4me3-BvCR in the DDR revealed that survivin-sensitive genes (n=28) gained an increase in H3K4me3 tags, while the IFNγ-sensitive genes (n=45) increased both H3K4me3 and H3K27me3 deposition after YM155 treatment (**Supplementary Figure S3D**).

Analyzing the transcriptional response mediated by IFNγ and survivin through H3K4me3-BvCR of the DDR network, we found that 11/45 IFNγ-sensitive genes (*FANCA, DPF2, RMI2, XRCC3, CDKN1A, PNKP, POLM, EXO5, MSH2, ATAD5, CUL4A*) (**Supplementary Figure S3C**) and 10/28 survivin-sensitive genes (**Figure 2G**) were annotated to one or more processes within the DDR network (*MGMT, STK11, MPG, XAB2, RFC1, FANCI, MSH6, MCM8, ATRX, ACTR2*). IFNγ signaling suppressed genes of the SSB/DSB repair and ubiquitin response (*FANCI, MSH6, MSH2, ATAD5*) and activated the genes involved in stress response and cell cycle control (*CDKN1A, STK11, RMI2, TP53, PIDD1*) (**Figure 2G, Supplementary Figure S3C**). Importantly, inhibition of survivin counteracted these effects of IFNγ and upregulated the DNA repair genes (**Figure 2G**), suggesting that a bimodal response to IFNγ treatment was survivin dependent.

Thus, interrogation of the DDR network controlled by H3K4me3-BvCR and survivin revealed that IFNγ stimulation activated the stress response and cell cycle control genes and repressed the DNA damage repair genes. Survivin acts as a critical mediator of IFNγ effects in regulating the DNA damage response processes mediated through H3K4me3-BvCR.

### 6. Binding of survivin to BvCR is sequence-specific and assembles TF complexes

Having detailed the functional significance of the survivin binding to BvCR in CD4+ cells, we investigated if nuclear TF known to read histone marks facilitated this interaction. Previous studies (Fukuda et al., 2015; Jensen et al., 2023; Erlandsson et al., 2022) reported that survivin’s control on transcription happened not in a solitary manner, but by association with transcription factors such as IRF1, SMAD3, and PRC2 complex. Hence, we performed a motif enrichment analysis of the DNA sequences in the survivin-positive (S+BvCR, n=4068) and survivin-negative BvCR (BvCR, n=2131), as well as in the survivin bound regions outside BvCR (Survivin, n=7823) (**Figure 3A**). Sequences ranging between 400 bp-700 bp were queried for motif presence and the highly enriched motifs (E-value>100) were annotated to the known human TF-binding motifs collected in the HOCOMOCO database. This analysis discovered that the S+BvCR and survivin peaks were frequently enriched with identical complex motifs (**Figure 3A**), which were estimated as potential binding sites of 331 TF within these genome regions (**Supplementary Table T3**). Notably, the TF motif enrichment was found solely in the survivin binding regions (S+BvCR and Survivin) (**Figure 3A**). The TF motif enrichment was absent or significantly lower in other BvCRs, which implied that survivin accounted for the sequence specificity of the binding.

**Figure 3.**
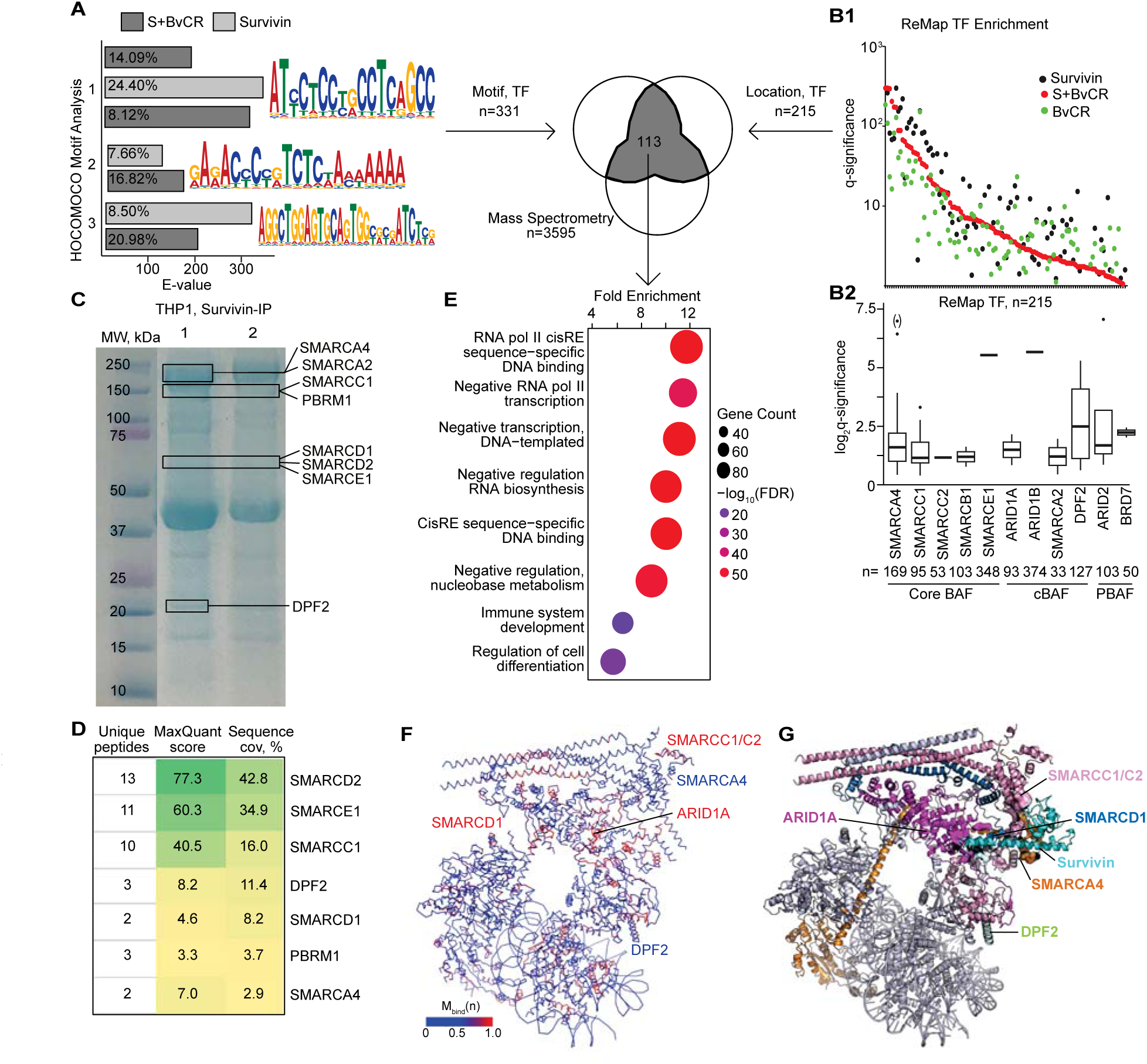
Survivin binding within H3K4me3-BvCR is sequence-specific and mobilizes BAF/SWI complex. (A) Bar plot of motif enrichment in survivin peaks within and outside of BvCR. MEME motifs enriched within BvCR. Venn diagram of TF identified by DNA sequence motif (MEME suite) and location (ChIP-seq) and mass spectrometry (MS). Motif enrichment within BvCR not containing survivin is not shown as no enrichment was observed. (B1) Dot plot of enrichment for human proteins/TF within BvCR, by ChIP-seq based ReMap2022 database. (B2) Box plot of enrichment for BAF/SWI complex proteins in ReMap2022 database. Counts underneath protein names indicates the number of overlaps with BvCR. (C) Coomassie-stained electrophoresis gel depicts survivin-bound proteins precipitated from THP1 cell lysate, separated by molecular weight (MW). Lanes represent two independent experiments (1 and 2, respectively). Bands with BAF/SWI complex proteins identified by mass spectrometry are indicated by boxes. (D) Table of BAF/SWI proteins identified by mass spectrometry. Enrichment score, number of unique peptides, and sequence coverage are calculated by MaxQuant software. (E) Bubble plot of biological processes enriched in proteins colocalized with survivin. Number of proteins in the process is indicated by the bubble size. Fold enrichment is shown by color intensity. (F) Ribbon diagram of the canonical BAF/SWI complex (PDB ID: 6LTJ) depicts the regions with a high probability of survivin binding (red) predicted by functional composition analysis. (G) Ribbon diagram depicting interaction between the human survivin-H3 tail complex (PDB ID: 3UEF) and the canonical BAF/SWI complex (PDB ID: 6LTJ) predicted by protein-protein docking analysis.

To explore this finding from another angle, we integrated the genomic location of BvCR, S+BvCR, and survivin peaks with the location occupied by human TFs identified via chromatin sequencing analysis in a diversity of human cells, available within the ReMap2022 database (Hammal et al., 2022). The TF enrichment analysis in the integrated datasets revealed that 215 TFs had a prevalent binding within BvCR (**Figure 3B, Supplementary Table T3**). The strongest TF enrichment was observed in the survivin binding regions (S+BvCR and Survivin), and the enrichment was weaker in BvCR not containing survivin (**Figure 3B1**). The cumulative analysis of the ChIP-seq datasets for the core subunits of the DDR pathway controlling BAF complex SMARCA4/BRG1, SMARCC1, SMARCC2, SMARCB1 and SMARCE1 demonstrated a large number of genomic overlaps with BvCR (**Figure 3B2**), which was visualized in THP1 cells for BRG1 earlier (**Figure 2E**). This finding suggests that survivin-BvCR binding mediates genomic colocalization and gene transcription (**Figure 2E**) of the BAF/SWI complex. Remarkably, the core BAF complex subunits common for cBAF and PBAF complexes overlapped frequently (∼40%) with H3K4me3-BvCR (**Supplementary Figure S5A**), while the integration of complex-specific subunits had a lower enrichment ratio with no preference for either cBAF or PBAF complexes (**Supplementary Figure S5A**). Similar enrichment was found for the cBAF subunits ARID1B, DPF2, SMARCA2 and PBAF subunits ARID2, and BRD7 (**Figure 3B2**).

Analysis of biological processes and molecular function revealed that the TF proteins identified by motif and genomic location analysis (n=113, **Supplementary Table T3**) were important in enacting sequence-specific binding of RNA Polymerase II to *cis*-regulatory elements (*cis*-RE) (GO:0000978) (**Figure 3E**). This validated our previous results that survivin and BvCR mediated control on gene transcription by situating within *cis*-RE of the human genome. Other chromatin-related processes such as RNA Polymerase II dependent transcription (GO:0000122) and negative RNA biosynthesis (GO:1902679) were also featured among the functions of TFs co-localized with survivin and BvCR (**Figure 3E**). Thus, the DNA sequence and location analyses of BvCR followed by biological pathway enrichment demonstrated that survivin binding assembled TF complexes, to support the regulation of gene transcription via *cis*-RE.

### 7. Survivin anchors BAF/SWI complex to BvCR

To experimentally develop chromatin sequencing results that indicated a survivin-BAF interaction in DDR function, we performed protein analysis of nuclear extract of THP1 cells immunoprecipitated (IP) with survivin. Survivin-IP material was separated by molecular weight using electrophoresis in two independent experiments, and proteomic analysis was done individually for each band by nano LC-ESI-MS mass spectrometry (MS) (**Figure 3C**).

The MS analysis revealed numerous peptides unique for the BAF complex components including SMARCA2/4, SMARCC1, SMARCC2, SMARCD1, SMARCD2, SMARCE1, DPF2, PBRM1 (**Figure 3C**). Several of the BAF complex proteins were reproduced in the second IP experiment (**Lane 2, Figure 3C**), suggesting a physical association of survivin with the BAF complex. Particularly, we observed high sequence coverage of the base module subunits SMARCC1, SMARCD1, SMARCE1, and SMARCD2 that anchored BAF complex to chromatin (**Figure 3D**).

To investigate in detail the interaction between survivin and the BAF complex, we applied the compositional analysis of proteins comprising the conventional (c)BAF and polybromo(P)BAF complexes aiming to predict their interaction with survivin. For this purpose, we utilized the peptide model based on the functional group composition of each protein of the complex (Jensen et al., 2023; Anindya et al., 2024). We identified that SMARCD1, SMARCA2, SMARCA4, SMARCC1, and BRD7 exhibit comparable ratios between the predicted binding peptides to the total number of generated peptides (R_bind_) values in the range of 0.39-0.43. These proteins harbor a high fraction of regions favorable to survivin, while SMARCE1 shows a lower R_bind_ of 0.32 (**Supplementary Table T5**). DPF2 and PHF10 show enrichment in survivin binding regions, with R_bind_ values of 0.46 and 0.51, respectively. Conversely, ARID1A and ARID2 exhibit limited regions with R_bind_ values of 0.24 and 0.19, respectively (**Supplementary Table T5**), compared to the median R_bind_ of the proteome. Our previous observations indicate that only specific domains of the core subunit of the PRC2 complex JARID2 contain a survivin binding region, despite large regions of JARID2 not binding survivin (Jensen et al., 2023). Therefore, mapping to sequence or 3D space becomes essential to identify clusters of polypeptide regions with suitable composition. As a result, we tested robustness of the binding by introducing mutations in the predicted binding positions of the peptides. The ratio between the number of mutations that support binding to the total number of mutations (M_bind_(n)), was mapped onto the 3D structure of cBAF and the sequence of its subunits (**Figure 3F, Supplementary Figure S5B).** The exposed predicted survivin binding regions are apparent as red-marked residues on the solvent accessible ribbon representation of cBAF.

Inspired by the results obtained by the peptide-based prediction, we conducted protein-protein docking between survivin and the subunits of the BAF complex. Structural modelling of the binding interphase between the BAF complex and survivin demonstrated that survivin exhibits maximal binding contacts with amino acid residues of SMARCC1, SMARCC2, SMARCD2, SMARCE1 (**Figure 3G**). The predicted complex structures of survivin binding with the cBAF and PBAF complexes are depicted in **Figure 3G** and the **Supplementary Figure S5C**, respectively. This analysis generates the hypothesis cBAF and PBAF complexes together with survivin involve close interactions with BRG1/SMARCA4, SMARCC1/C2, and SMARCD1 subunits. The absence of interaction between SMARCE1 and survivin in our predicted complexes (**Supplementary Figure S5B**, **Supplementary Table T5**) may be attributed to the incompleteness of the electron microscopy structures available for the cBAF (only residue range 172-276 is visible) and PBAF complexes. Additionally, cBAF-specific subunits ARID1A, and PBAF-specific subunits PHF10, contributed to the predicted interactions with survivin, while the amino acid residues of DPF2, ARID2, PBRM1, and BRD7 subunits had predicted interaction with survivin in the docking simulations, but their composition compatibility with survivin (M_bind_(n)) in the docking interface is low. These docking analyses highlighted the distinctive roles of specific SWI subunits in chromatin accessibility regulation (Mashtalir et al., 2018; Ho et al., 2009). Intriguingly, the cBAF subunits exhibited a larger interaction area with survivin compared to the PBAF subunits (34 residues from cBAF vs. 21 residues from PBAF) and engaged the residues in slightly different protein interfaces (**Supplementary Figure S5D**). Accordingly, the docking energy score for the survivin-cBAF and survivin-PBAF complexes was -363.7 and -318.6 kcal/mol, respectively. These results present a structural model of survivin binding with BAF/SWI complex through the DNA anchoring module and suggest a slight preference of binding to the cBAF complex over the PBAF complex. The conceived interaction between survivin and BAF complex still awaits experimental proof of physical contact in the predicted protein-protein interfaces.

Taken together, the combination of biomolecular interaction experiments through mass spectrometry, compositional analysis and structural modelling, advocates in favor of survivin aiding the anchoring of BAF complexes to BvCR.

### 8. IFNγ affected cell cycle progress in CD4^+^ cells, while inhibition of IFNγ signaling and survivin caused cell cycle arrest and the DNA damage recognition

Integrating all the results presented above, we propose a functional model according to which the DDR network in CD4^+^ cells is controlled by survivin-positive BvCR located within *cis*-RE of the human genome (**Figure 4A**). To aid this function, survivin anchors the BAF complex to the BvCR via transduction of the activating effects of IFNγ signaling in these cells. We sought to gather experimental verification of this model *in vitro* in IFNγ-stimulated CD4^+^ cells.

**Figure 4.**
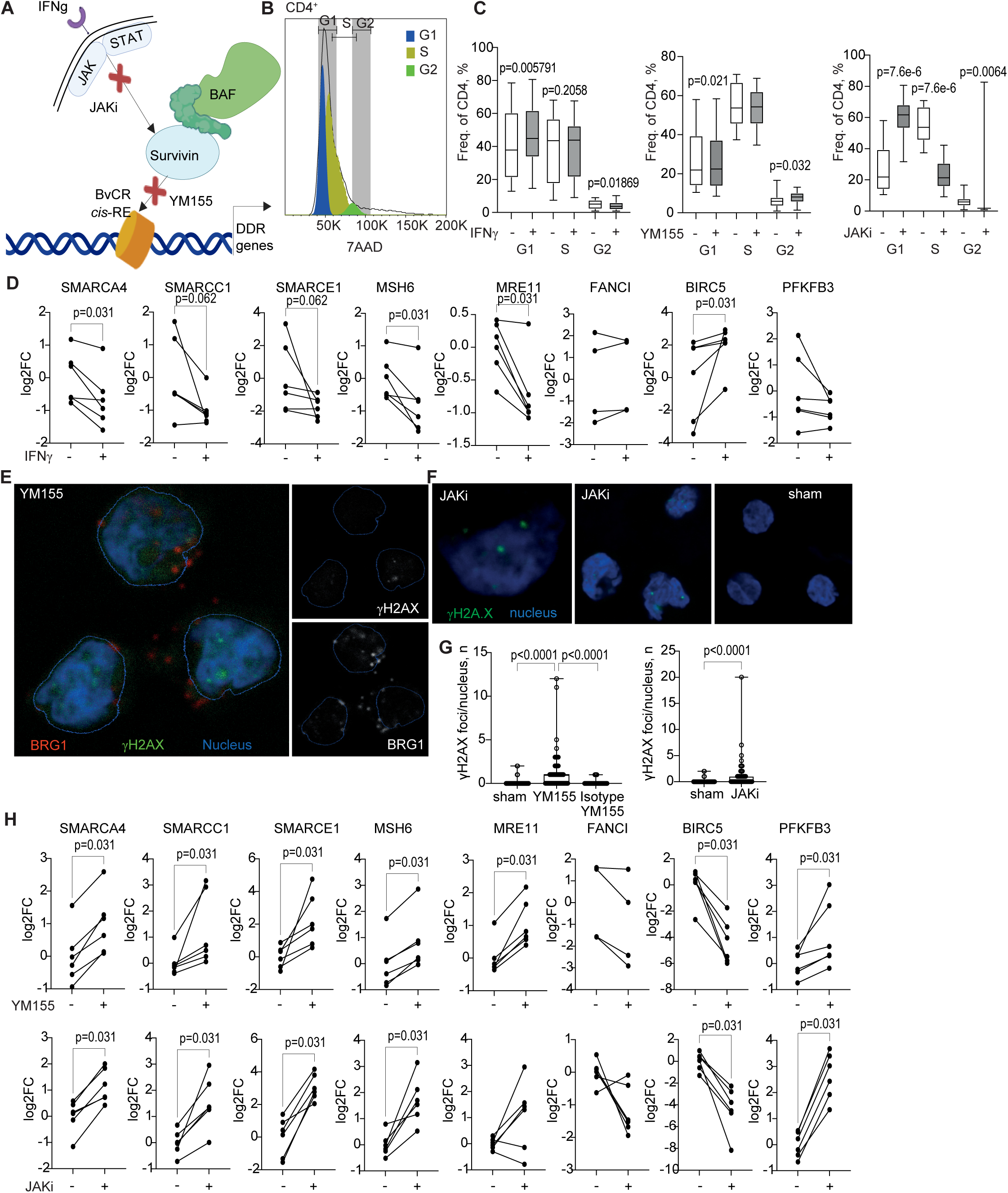
Survivin inhibition and JAKi treatment enhances DNA damage repair and disrupts cell cycle progression. (A) A model of intervention in IFNγ, survivin-BAF complex and BvCR dependent transcription of DNA damage response (DDR) genes by treatment with JAK-inhibitor (JAKi) and survivin inhibitor YM155. Cis-RE, regulatory element. (B) Histogram of 7AAD^+^CD4^+^ cell distribution by phases of the cell cycle. Colored areas indicate G1 (blue), S (yellow) and G2 (green) phases. (C) Frequency distribution of 7AAD+ cells by cell cycle phases in CD4^+^ cells treated with IFNγ (50ng/ml), YM155 (10nM), and JAKi (10µM) compared to sham (DMSO). P-values are obtained by Wilcoxon paired test. (D) Linear plot of gene expression change in CD4^+^ cells treated with IFNγ (50ng/ml for 48h) measured by qPCR. Expression difference was calculated by Wilcoxon paired test. (E, F) THP1 cells were treated for 24h with YM155 (200nM), JAKi (50 µM), or sham (DMSO), fixed and stained for DNA damage using antibodies against BRG1 (red), γH2AX (green) and nuclear stain (blue). (G) γH2AX foci in nuclei were counted in each nucleus of 90-200 cells per treatment using ImageJ. P-values are obtained by Mann-Whitney U test. (H) Linear plot of gene expression change in CD4^+^ cells treated with YM155 (10nM) or JAKi (10 μM) for 48h, compared to sham (DMSO), measured by qPCR. Expression difference was calculated by Wilcoxon paired test.

Since cell cycle featured among the pathways controlled by the BvCR (**Figure 2C**), we investigated the cell cycle progress in CD4^+^ cells using the DNA binding fluorescent dye 7-aminoactinomycin D (7AAD) in flow cytometry (**Figure 4B)**. We observed that the progress of cell cycle from G1 to S and further to the G2 phase was significantly accelerated by IFNγ treatment (**Figure 4C**). The YM155-treated cells slowed down progression of cell cycle causing accumulation of cells in the G1 and G2 phases (**Figure 4C**). Inhibitors of JAK/STAT pathway (JAKi) downstream of IFNγ receptor induced cell cycle arrest in G1 phase, which depleted the cultures for cells in G2 phase (**Figure 4C**). Therefore, both JAKi treatment and survivin inhibition disrupted proper cell cycle progression in CD4^+^ cells.

Expectedly, IFNγ stimulation activated the functional link between survivin and energy supply in CD4^+^ cells by upregulating *BIRC5* and repressing *PFKFB3* (**Figure 4D**) (Erlandsson et al., 2022), a gene that is important for efficient homologous recombination (Gustafsson et al., 2018). Notably, the IFNγ-treated cells presented repression of the core BAF complex genes *SMARCA4*, *SMARCC1*, *SMARCE1* (**Figure 4C**) and had significantly lower transcription of the central DNA repair genes *MRE11*, *MSH6*, and *FANCI* connected to the H3K4me3-BvCR in CD4^+^ cells (**Figure 4C**). Survivin inhibition by YM155 promoted the accumulation of γH2AX^+^ foci in THP1 cells (**Figure 4E, G**). A similar accumulation of γH2AX foci was observed after JAKi treatment (**Figure 4F, G**). The cells treated with JAKi and YM155 had significantly suppressed transcription of the survivin gene *BIRC5,* which was followed by upregulation of the BAF complex genes and the DNA damage repair genes (**Figure 4H**) in consistency with the RNA sequencing results (**Figure 2G)** and accumulation of the damaged DNA marked by γH2AX (**Figure 4G**). This finding indicates that survivin-BAF axis connects IFNγ signaling to DDR through survivin but has a suppressive effect on core BAF complex transcription.

Taken together, the *in vitro* studies validated the proposed functional model which states that survivin exploits bivalency using IFNγ signaling in CD4^+^ cells, aiding in connecting the BAF complex to its DDR activity. Inhibition of IFNγ signaling and survivin was associated with cell cycle arrest and accumulation of damaged DNA. This triggered the subsequent activation of DNA repair genes, including transcription of the BAF complex.

### 9. BRG1 expression defined a specific phenotype of CD4^+^ cells in patients with rheumatoid arthritis

To investigate the requirement of the BAF/SWI complex in mediating the survivin-dependent DNA damage control in autoimmune CD4^+^ cells, we used transcriptome data sets of CD4^+^ cells of 24 RA patients (GSE190349). Guided by BRG1*/SMARCA4* transcription, we found that both survivin and IFNγ were highly co-expressed in the *BRG1^hi^*CD4^+^ cells (**Figure 5A).** Mapping of the differentially expressed genes to the DDR network revealed that the nodes of DNA repair, replication, and G1 arrest were upregulated in BRG1^hi^ cells (**Supplementary Figure S6B**), suggesting unbalanced control of the DDR in these cells.

**Figure 5.**
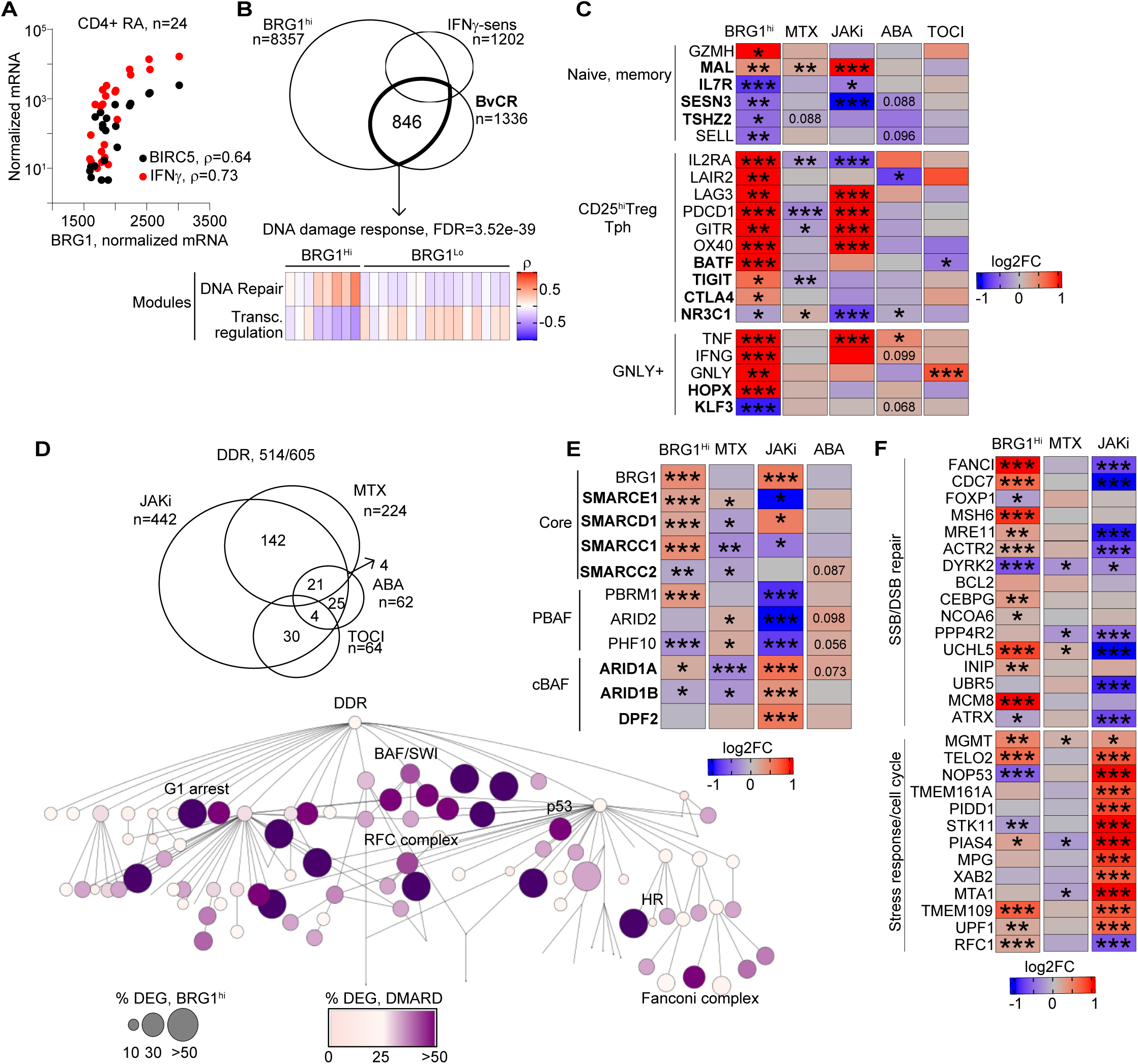
Immunomodulating treatment affects DNA damage response in CD4^+^cells of patients with rheumatoid arthritis. (A) Dot correlation plot of normalized mean expression of *BRG1, BIRC5* and *IFNG* genes in CD4^+^ cells of patients with rheumatoid arthritis. Spearman’s rho values are indicated. (B) Venn diagram of differentially expressed genes (DEG) in BRG1^hi^CD4^+^ cells connected to BvCR and in IFNγ-treated CD4^+^ cells. Heatmap of Spearman’s rho correlation values of genes connected to BvCR in BRG1^hi^ and BRG1^lo^ cells identified by weighted correlation network analysis (WGCNA). (C) Heatmap of expression difference in T cell specific markers identified by RNA-seq in BRG1^hi^CD4^+^ cells and in CD4^+^ cells before and after treatment with abatacept (ABAT, n=14), tocilizumab (TOCI, n=6) and methotrexate (MTX, n=28) and in CD4^+^ cells of JAKi-treated (n=23) and untreated (n=9) RA patients. Expression difference in CD4^+^ cells before and after treatment (for ABAT, TOCI and MTX) and in JAKi treated and untreated patients was calculated by DESeq2. Nominal p-values are indicated. * < 0.05, ** < 0.01, *** <0.001. (D) Venn diagram of DEG changed with treatment in the DNA damage response (DDR) network. DDR network map of DEG changed with treatment. Node size indicates the percentage of BRG1^hi^ DEG. Node color indicates the percentage of DEG in the node. (E) Heatmap of expression difference in BAF/SWI complex proteins in BRG1^hi^CD4^+^ cells and in CD4^+^ cells after treatment, by RNA-seq. Expression difference was calculated by DESeq2. Nominal p-values are indicated. * < 0.05, ** < 0.01, *** <0.001. (F) Heatmap of expression difference in DDR network genes in BRG1^hi^CD4^+^ cells and in CD4^+^ cells after treatment.

Next, we investigated if genes connected to BvCR were abnormally regulated in BRG1^hi^ cells and if it was associated with the pathogenic phenotype in RA CD4^+^ cells. A total of 63% (846/1336) of the BvCR connected genes involved in at least one of the main pathways controlled by BvCR were differentially expressed in BRG1^hi^CD4^+^ cells in RA patients, including the DDR pathway (**Figure 5B**). To identify the functional modules of the genes co-expressed with BRG1 in the BRG1^hi^CD4^+^ cells, we applied the weighted correlation network analysis (WGCNA), asking if the hub genes that showed high co-expression in the BRG1^hi^ and BRG1^lo^ cells were functional in the biological processes that had been identified experimentally.

The WGCNA approach identified two modules, where the hub genes were positively associated with BRG1^hi^ cells (n=498) and with BRG1^lo^ cells (n=348) (**Figure 5B, Supplementary Table T4**). Several core subunits of the BAF complex including BRG1/*SMARCA4, SMARCC1, SMARCD1, SMARCE1* and cBAF specific subunit *ARID1A* were accumulated in the module of BRG1^hi^ cells (**Figure 5E**). In addition to the cBAF complex subunits, this module showed the enrichment for the DNA repair pathway (GO:0006281, FDR=2.8e-34) and cell cycle pathway (R-HSA-1640170, FDR=1.3e-36) (**Figure 5B**). Common cell cycle genes *TP53 and CDKN1A* controlled by BvCR were also highly upregulated in the *BRG1^hi^* cells while the *ATM* gene was repressed. The module of BRG1^hi^ cells included the IFNγ-sensitive genes of *PIAS4*, *MSH6*, *FANCI*, *MRE11, TELO2*, and *SMC3* (**Figure 5F**) that were previously shown to change H3K4me3 tag deposition in the connected BvCR after survivin inhibition (**Figure 2F, Supplementary Figure S4**). In contrast, several regulators of transcription were present in the module of BRG1^lo^ cells (FDR=1.35e-70) including *FOXP1, CBX4, PBX1, LEF1*.

To translate the profile of BRG1^hi^CD4^+^ cells into joint pathology in rheumatoid arthritis, we utilized characteristics of the inflammatory cell subsets identified in RA synovia by simultaneous quantification of cell surface proteins and transcriptomic data within a single cell readout (Zhang et al., 2023). Of the 24 T cell clusters, we focused only on CD4^+^ and investigated if genes of these clusters were differentially expressed in the BRG1^hi^CD4^+^ cells and connected to BvCR. Based on the key markers of the synovial clusters, we found that transcriptome profiling of *BRG1^hi^*CD4^+^ cells was enriched in the synovial cytotoxic *GNLY^hi^HOPX^hi^* cells expressing *IFNG, TNF* and *GZMA*, and the peripheral T helper cells abundant in the immune check-point receptors PD1*/PDCD1*, *CTLA4*, and *LAG3* (**Figure 5C**). These pathogenic T cells expressed high levels of OX40/*TNFRSF4*, GITR/*TNFRSF18*, *LAIR2*, common with the CD25^hi^T cell subset, and combined the ability to infiltrate inflamed tissue with the ability to drive autoimmunity in RA (Okazaki et al., 2013; Rao et al., 2017; Kostine et al., 2018). In contrast, the naïve *SELL*/CD62L^hi^ and memory T cell subsets recognized by *IL7R, SESN3, KLF3, NR3C1* and *TSHZ2* genes connected to the BvCR were under-represented in BRG1^hi^CD4^+^ cells. Notably, the genes of about half of the pathogenic subset markers were controlled by BvCR and sensitive to IFNγ (**Figure 5C**).

Taken together, the deregulated control of the DDR pathway in BRG1^hi^CD4^+^ cells was demonstrated here as the feature of the pathogenic peripheral T helper cells abundant in RA synovia and contributing to their pathology.

### 10. Effect of immunomodulating treatment on the BvCR dependent profile of the pathogenic BRG1^hi^CD4^+^ cells

Having characterized the pathogenic profile of the BRG1^hi^CD4^+^T cells of patients with rheumatoid arthritis, we asked how immunomodulating treatment influenced the DDR network in CD4^+^T cells. To investigate this, we analyzed the paired transcriptome data of CD4^+^ cells obtained before and after treatment with CTLA4 fusion protein abatacept (n=14, GSE121827; Glatigny et al., 2019), IL6 receptor blocking antibody tocilizumab (n=6, GSE113156, Takeshita et al., 2019), and methotrexate (n=28, GSE176440; Shoda et al., 2022). Additionally, the effect of JAKi on CD4^+^ cells was studied using cross-sectional transcriptome of JAKi-treated (n=24) in comparison with untreated RA patients (n=9, GSE201669). Demographic characteristics of the patients are shown in **Supplementary Figure S6A.**

A total of 514 of 605 genes annotated to the DDR network were responsive to the immunomodulating treatment (**Figure 5D**). The predominant part (442/605, 73%) of the DDR network genes were changed in the JAKi-treated patients. Methotrexate treatment accounted for more than one-third of the differentially expressed genes (224/605, 37%) followed by abatacept and tocilizumab in equal proportions **(Figure 5D).** Interestingly, a large portion (>50%) of the DEG in BRG1^hi^ cells that were functional in the DDR network processes of BAF/SWI complex, G1 arrest, p53 pathway, DNA replication by RFC complex, and Fanconi complex, were responsive to at least one of the treatments (**Figure 5D**). The genes of SSB/DSB repair and stress response, in which we had earlier detailed the effect of survivin and H3K4me3-BvCR (**Figure 2G**, **Figure 4H**), the effects of methotrexate and JAKi frequently opposed each other (**Figure 5F**). Methotrexate changed the expression of SSB/DSB repair genes *DYRK2*, *PPP4R2* and *UCHL5* and the stress response genes *MGMT*, *MTA1*, *PIAS4* (**Figure 5F**). Interestingly, JAKi treatment upregulated all stress response genes and repressed the DNA repair genes (**Figure 5F**). The effect was pronounced in the specific genes of *FANCI, CDC7, MCM8, MGMT,* and *MRE11* which reversed the expression observed in BRG1^hi^ cells, suggesting that JAKi intervened in the BvCR-controlled DDR in CD4^+^ cells. We did not observe any transcription change in the important SSB mismatch repair genes *MSH6* and *MSH2* after methotrexate or JAKi treatment, suggesting that the effects on these genes observed in CD4^+^ cultures leveled off with longer drug exposure.

Since different BAF complexes have been reported to function in specific cellular processes (Begg et al., 2024; Ho et al., 2009), we analyzed the differential expression of cBAF and PBAF complex subunits after the immunomodulating treatment. Intriguingly, methotrexate treatment upregulated PBAF-specific subunits *ARID2* and *PHF10*, and downregulated cBAF subunits *ARID1A* and *ARID1B* (**Figure 5E**). In contrast, JAKi treatment upregulated BRG1 and cBAF subunits *ARID1A*, *ARID1B*, and *DPF2*, and downregulated PBAF subunits *PBRM1*, *ARID2*, and *PHF10* (**Figure 5E**). The effect of methotrexate on PBAF subunits was reciprocal with that of IFNγ, while JAKi suppressed PBAF probably independently of IFNγ due to their low IFNγ sensitivity (**Figure 5E**). Abatacept tended to upregulate both *PHF10* and *ARID1A*, while tocilizumab did not have any effect on either of the BAF/SWI complex genes. The core subunits *SMARCC1* and *SMARCD1*, which had been predicted in this study as the survivin interactors anchoring the BAF complex to H3K4me3-BvCR (**Figure 3**), were downregulated by both methotrexate and JAKi. The relative deficiency in these subunits will impair BAF complex function in recognition of DNA damage (Harrod et al., 2020; Begg et al., 2024) leading to accumulation of γH2AX on chromatin as demonstrated in *in vitro* experiments (**Figure 4**). These findings suggest that RA treatment differentially affected the processes within CD4^+^ cells potentially impairing BAF complex composition and its mobilization to the DNA damage sites.

Taken together, immunomodulating treatment altered transcription of the BvCR-connected genes which were active within the processes of DNA repair and stress response. The treatment dependent modification in composition of the BAF complex could explain a loosened affinity of the complex to BvCR, which reduced the transcriptional control.

## Discussion

In this study, we uncover the function of BvCR in orchestrating transcriptional response to DNA damage in adaptive immunity. Survivin is exposed here as an essential facilitator in this process by ensuring an appropriate balance of the histone H3 epigenetic marks deposition in the BvCR. We demonstrated a tight link between the survivin-dependent H3 marks deposition in BvCR in human CD4^+^ cells and activation of DNA damage response genes connected to these BvCR. Through sequence analysis of survivin-bound BvCR and structural modelling of interactions between survivin and BAF complex, we revealed that survivin countered IFN-mediated effects of DNA damage signaling and hampers DNA repair activity. In autoimmune CD4^+^ cells of patients with rheumatoid arthritis, we identified the pathogenic profile of the BAF complex expressing cell subset. Furthermore, we demonstrated that the BAF complex proteins are targeted by existing immunomodulating anti-rheumatic drugs which was associated with a change in the pathogenic profile of the cells.

Our study revealed the importance of BvCR in finetuning transcriptional characteristics of distal *cis*-RE of CD4^+^ T cells, which shed new light on BvCR function in mature T cells. This is a valuable addition to the known role of BvCR in resolving T cell lineage choice (Wei et al., 2009; Roh et al., 2006; Kinkley et al., 2016). Survivin sought binding to BvCR by specificity to DNA sequence, opening a platform for TF cooperation that read H3 marks in the BvCR (Jensen et al., 2023). Through the occupancy of BvCR, survivin balanced the quantitative level of H3K4me3 and H3K27me3 deposition in cis-RE with preferable recognition of the BvCR dominated by H3K4me3. The phenomenon of a dominant H3 mark within BvCR has been suggested to provide insight into functional cell phenotypes, e.g., control of the immune system by H3K4me3-dominant and control of development by H3K27me3-dominant BvCR (Kinkley et al., 2016). Such epigenetic classification was also appealing clinically for computational dissection for treatment choice in cancer (Alarcón et al., 2021).

Further, we propose that survivin binding to the H3K4me3-BvCR facilitates a sequence-specific anchoring of the BAF/SWI complex. Indeed, the BAF/SWI complex is attracted to specific genomic locations through their accessory subunits and facilitates chromatin remodeling by removal and replacement of histone variants (Harrod et al., 2020; Clapier et al., 2017). In support of this, we observed a significant change of histone H3 marks after survivin inhibition and molecular interactions between survivin and the base of the BAF/SWI complex including subunits SMARCC1, SMARCB1, SMARCD1, and SMARCE1 (He et al., 2020), which illustrated how BAF/SWI complex was mobilized to chromatin. This model finds validation in a recent functional study which deduced the role of epigenetic modifications for affinity of BAF complex binding to chromatin (Mashtalir et al., 2021), and in our recent study which showed that survivin inhibited the PRC2 repressive complex which obstructed transcription (Jensen et al., 2023). Thus, the proposed interaction mode of survivin with the BAF/SWI complex combined with its inhibitory action on the PRC2 complex potentially permits gene transcription. BAF/SWI and PRC2 complexes display antagonistic and mutually exclusive binding at the same locus (Leitner et al., 2020; Januario et al., 2017). By adopting mutually exclusive modes of interaction, survivin co-opts the bivalent genome’s modulation of DNA damage response or transcriptional repression, respectively. The contrasting modes of survivin interaction with the BAF/SWI and PRC2 complexes enhance survivin-mediated control on genomic bivalency.

BAF/SWI complex has been shown to regulate expression of cytokine genes (Jeong et al., 2010), suggesting a connection between genome organization and immune signaling pathways. Aberrant genome organization activates IFN signaling through an increased DNA damage response, identified as one of the emerging functions of the BAF complex (Harrod et al., 2020; Begg et al., 2024; Ribeiro-Silva et al., 2019). Our study shows that survivin is an intermediary in this connection and propagates IFN-mediated effects in CD4^+^T cells through accelerating cell cycle progression. Survivin inhibition as well as inhibition of JAK/STAT signaling counteracts the IFN-induced effect through the cell cycle arrest and upregulation of DNA repair genes. Notably, survivin binding to BvCR rendered transcriptional control of the cBAF complex subunits *ARID1A, ARID1B, DPF2*, and SS18, but not the PBAF complex. Together with the shown association of survivin with the cBAF complex, our data support the view that survivin adjustment of BAF complex composition by localizing with BvCR help in the efficiency of DNA repair activity. This proposal is in line with a recent structural report suggesting that cBAF enables chromatin accessibility by occupying distal enhancers (Otto et al., 2023; Begg et al., 2024). Additionally, these findings build on our previous study which has shown that survivin transcriptionally controls *PFKFB3* (Erlandsson et al., 2022), the glycolytic enzyme which plays a crucial role in promoting DNA repair (Gustafsson et al., 2018) and encouraging metabolic adaptation in IFN-producing CD4^+^T cells. Together, our data exposed the DNA damage signaling as one of IFN-dependent processes in CD4^+^T cells coordinated by survivin and the conventional BAF/SWI complex.

This study reveals an unappreciated role of BAF complex function in the context of autoimmunity. While BAF/SWI complex subunits are known to be mutated in about 20% of cancers (Kadoch et al., 2013), their role in autoimmune conditions remains unknown. We found that cBAF activity dominates in BRG1^hi^CD4^+^ cells of RA patients. In these cells, the high survivin and cBAF complex levels coexisted with upregulated DNA repair profiles. The BRG1^hi^CD4^+^ cells were highly enriched in the immune checkpoint and other receptors characteristic for the peripheral T-helper CD4^+^ cells identified in the RA synovial tissue (Zhang et al., 2023). This similarity outlines the pathogenic potential of the BRG1^hi^CD4^+^ cells capable of homing to the joint to induce and propagate arthritis.

Evaluation of intervention in the DNA damage response pathway induced by immunomodulating RA treatment affected the transcriptional effects of survivin-containing BvCR in assisting IFN signaling in CD4^+^T cells. Activation of DNA damage response is a crucial characteristic of autoimmune CD4^+^T cells, which accumulate impaired DNA repair, induction of apoptosis, and abnormal metabolism (Shao, 2018; Manolakou et al., 2021). Methotrexate treatment mildly diverged DNA damage response but JAKi strongly upregulated DNA damage signaling, together suggesting that immunomodulating treatment intervene in these IFN-dependent processes in CD4^+^ cells. More importantly, we observed an adjustment in BAF/SWI complex composition, where methotrexate or JAKi treatment differentially upregulated the BAF complex subunits to enact DNA damage response. However, both drugs downregulated the base module subunits *SMARCE1, SMARCD1, SMARCC2,* where survivin potentially acts as an intermediary mobilizing BAF/SWI complex to BvCR. Thus, different treatment modalities mitigate survivin mobilization of the BAF complex to the BvCR and elicit a shift in composition of the BAF complex. Together, these processes modulate the DNA damage signaling activity in autoimmune CD4^+^ cells alleviating arthritis pathology.

In conclusion, our study evidenced a hitherto undefined connection between BvCR and survivin in effector CD4^+^T cells to assure the promotion of IFNγ-induced DNA damage response ubiquitous in autoimmunity. DNA repair and stress response were both activated in BRG1^hi^BIRC5^hi^ arthritogenic CD4^+^ cells. Diametric regulation by either of the mechanisms after treatment could possibly be explained by composition of the BAF complex prevalent under given conditions. Intervention with the BAF complex mobilization to the bivalent chromatin mitigating survivin function, offers a new platform for drug development targeting autoimmunity.

## Materials and methods

### Human material

Blood samples of 67 RA patients and 40 healthy controls, all collected at the Rheumatology Clinic, Sahlgrenska Hospital, Gothenburg, were used in this study. Clinical characteristics of the patients are shown in **Supplementary Figure S6A**. All RA patients fulfilled the EULAR/ACR classification criteria (Aletaha et al., 2010) and gave written informed consent before the blood sampling. The study was approved by the Swedish Ethical Review Authority (659-2011) and done in accordance with the Declaration of Helsinki. CD4^+^ cells from healthy controls were used for ChIP-seq, RNA-seq and qPCR after treatment with IFNγ, IFNγ+YM155, and JAK-inhibitor tofacitinib, and flow cytometry.

### Isolation and stimulation of CD4^+^ cells

Human peripheral blood mononuclear cells were isolated from the venous peripheral blood by density gradient separation on Lymphoprep (Axis-Shield PoC As, Dundee, Scotland). CD4^+^ cells were isolated by positive selection (Invitrogen, 11331D), and cultured at density 1.25×10^6^ cells/ml in wells coated with anti-CD3 antibody (0.5 μg/ml; OKT3, Sigma-Aldrich, Saint Luis, Missouri, USA), in RPMI medium (Gibco, Waltham, Massachusetts, USA) supplemented with 50 μM β-mercaptoethanol (Gibco), Glutamax 2 mM (Gibco), gentamicin 50 μg/ml (Sanofi-Aventis, Paris, France) and 10% fetal bovine serum (Sigma-Aldrich) at 37°C in a humidified 5% CO_2_ atmosphere. For RNA-seq, CD4^+^ cell cultures were treated with recombinant IFNγ (50 ng/ml; Peprotech, Cranbury, NJ, USA) and survivin inhibitor sepantronium bromide (Nakahara et al., 2007) (10 nM YM155, S1130, Selleck Chemicals, Houston, TX) for 72h. For qPCR, CD4^+^ cell cultures were treated with IFNγ, YM155, or JAK-inhibitor tofacitinib (10 µM; S2789, Selleck Chemicals) for 48h.

### DNA content and cell cycle analysis

Freshly isolated PBMC from 16 healthy persons were stimulated for 48h with anti-CD3 antibodies with tofacitinib (0 or 10µM) or YM155 (0 or 10nM). Eight cultures were additionally treated with recombinant IFNγ (0 or 50 ng/ml) for the last 18 hours. At harvest, supernatants were collected, and cells were first blocked with Fc-block (BD 564220), stained with AF647-conjugated antibodies to CD4 (Biolegend 317422) followed by permeabilization (BD cytofix/cytoperm), and incubated overnight, 4°C, with 20 μg/ml of 7-aminoactinomycin D (7AAD, Invitrogen A1310) in perm/wash (BD Biosciences), and resuspended in 200 μl FACS buffer. Cells were acquired with flow cytometry system BD FACSLyric^TM^ (BD Biosciences) and data analysis was performed in the Tree Star FlowJo software using the inbuilt cell cycle analysis tool, Watson model with constrains, CV (G2) = CV (G1).

### Conventional quantitative (q)PCR

RNA was isolated with the Total RNA Purification Kit (#17200, Norgen Biotek). RNA concentration and quality were evaluated with a NanoDrop spectrophotometer (ThermoFisher Scientific) and Experion electrophoresis system (Bio-Rad Laboratories). cDNA was synthesized from RNA (400 ng) with the High-Capacity cDNA Reverse Transcription Kit (Applied Biosystems, Foster City, CA, USA). Real-time amplification was done with RT2 SYBR Green qPCR Mastermix (Qiagen) and a ViiA 7 Real-Time PCR System (ThermoFisher Scientific) as described (Andersson et al., 2017).

#### Primer Design

Primers were designed in-house using the Primer3 web client (https://primer3.ut.ee/). When applicable, primers were separated by an exon-exon boundary. Amplicon and primer size was limited to 60-150 and 18-24 base pairs, respectively. Melting temperature was set between 60-63°C, max poly-X to 3 and GC-content was limited to 40-60%. Primers suggested by the software was checked in Net Primer web client (https://www.premierbiosoft.com/netprimer/) for possible hairpin and primers-dimer structures. Lastly, correct binding of primers was validated in UCSC In-Silico PCR web client (http://genome.ucsc.edu/cgi-bin/hgPcr) against the GRCh38/hg38 human genome assembly. Primers used in this study are shown in **Supplementary Figure S6C**. Expression was calculated by the ddCt method.

#### Transcriptional sequencing (RNA-seq)

RNA from the CD4^+^ cell cultures was prepared using the Norgen Total RNA kit (17200 Norgen Biotek, Ontario, Canada). Quality control was done by Bioanalyzer RNA6000 Pico on Agilent2100 (Agilent, Santa Clara, CA, USA). Deep sequencing was done by RNA-seq (Hiseq2000, Illumina) at the core facility for Bioinformatics and Expression Analysis (Karolinska Institute, Huddinge, Sweden). Raw sequence data were obtained in Bcl-files and converted into fastq text format using the bcl2fastq program from Illumina.

#### Chromatin immunoprecipitation and sequencing (ChIP-seq)

For survivin-ChIP-seq analysis, CD4^+^ cells isolated from 12 women were stimulated with concanavalin A (ConA, 0.625 μg/ml, MP Biomedicals), and lipopolysaccharide (LPS) (5 μg/ml, Sigma-Aldrich) for 72 h (the samples were pooled in 4 independent samples after DNA purification). For histone-ChIP-seq analysis, CD4^+^ cells isolated from 3 women were stimulated with ConA and LPS as above for 24 h and then treated with YM155, 0 or 10 ng/ml, for 24 h, (the samples were pooled after DNA isolation). The cells were cross-linked and lysed with the EpiTect ChIP OneDay kit (Qiagen 334471), as recommended by the manufacturer. After sonication to shear the chromatin, cellular debris was removed by pelleting. After preclearing, 1% of the sample was saved as an input fraction and used as background for deduction of nonspecific chromatin binding. Pre-cleared chromatin was incubated with 2μg of anti-survivin (10811, Santa Cruz Biotechnology, Santa Cruz, CA, USA), anti-H3K27ac (C15410196, Diagenode), or anti-H3K27me3 (C15410195, Diagenode) or anti-H3K4me3 (C15410003, Diagenode). The immune complexes were washed, the cross-links were reversed, and the DNA was purified with the EpiTect ChIP OneDay kit (Qiagen) as recommended by the manufacturer. The quality of purified DNA was assessed with TapeStation (Agilent, Santa Clara, CA, USA). DNA libraries were prepared with ThruPLEX (Rubicon) and sequenced with a Hiseq2000 sequencing system (Illumina) according to the manufacturer’s protocols. Bcl-files were converted and demultiplexed to fastq with bcl2fastq (Illumina).

#### Immunohistochemistry and imaging

Human monocytic cell line THP1 (TIB-202, ATCC, Manassas, VA, USA) were seeded 10^6^/ml on glass chamber slides (Thermo Scientific) precoated with poly-L-lysine (Sigma-Aldrich, Saint Louis, MO, USA). Cells were treated with 10µM or 50µM tofacitinib or 200nM YM155 (both from Selleck chemicals) for 24h. At harvest, cells were fixed with 4% buffer and formalin for 10 minutes and blocked and permeabilised for 3 hours with 3% normal goat serum and 1% TritonX100. Primary antibodies against Histone H2A.X phosphorylated at Ser^139^ (γH2AX, mouse, Millipore 05-636), BRG1/SMARCA4 (rabbit, Bethyl Laboratories A300-813A), H3K4me3 (rabbit, C15410003, Diagenode), and PE-conjugated survivin (mouse, Clone 91630, RnD systems) and isotype controls were diluted in blocking buffer and the slides were incubated over night at 4°C. This was followed by Alexa-fluor conjugated secondary antibodies donkey-anti-mouse AF488 (Invitrogen A-21202), goat-anti-rabbit AF488 (Invitrogen A11034) or donkey-anti-rabbit AF647 (Invitrogen A-31573) for 2 hours at room temperature. Autofluorescence was blocked with 0.5% Sudan Black B (Sigma-Aldrich) in 70% ethanol for 20 min at room temperature. Nuclei were stained with Hoechst 34580 (NucBlue Live Cell Stain; Thermo Fisher Scientific) for 20 min and mounted with ProLong Gold antifading mounting reagent (Invitrogen).

#### Confocal imaging and analysis

Fluoresence microscopy was performed using the confocal imaging system Leica SP8 (Leica Microsystems, Wetzlar, Germany) with sequencial acquisition using a 40x oil objective and up to 10x digital zoom. The images were aquired at high resolution (1.5x digital zoom) viewing 40-70 nuclei per image. Within each sample, gH2AX-positive foci were enumerated in 2-3 images resulting in 78 to 222 nuclei per treatment.

Images were analysed with ImageJ version 2.9. Images were analysed in 8-bit composite images with threshold adjusted individually for each antibody to optimise the software’s ability to identify positive spots. Regions of interest (ROI, nuclei) were identified in the blue image (Hoechsts) using adjust threshold and analyze particles. Number of γH2AX-psotive foci in each nuclei was estimated in the green image (AF488 stain) within each ROI. Number of green foci per nuclus was estimated after noice reduction by despeckling using the ImageJ feature Find Maxima.

#### Affinity immunoprecipitation

THP-1 cells were cultured at a density of 3-10×10^5^ cells/mL in RPMI1640 medium supplemented with 10% FBS at 37°C in a humidified atmosphere of 5% CO2. Cells were lysed in modified RIPA-buffer (25 mM Tris-HCl pH 7.4, 200 mM NaCl, 1 mM EDTA, 1% NonidetP-40, 5% glycerol) supplemented with protease inhibitors (Complete mini, Roche), and immunoprecipitation was performed with 2μg of anti-survivin antibodies (RnD AF886) coupled to the Dynabeads Protein G Immunoprecipitation Kit (10007D, ThermoFisher Scientific), cross-linked with bis(sulfosuccinimidyl)suberate (A39266, Pierce™). The immune-precipitated complexes were washed extensively with the provided washing buffer plus fragment stream buffer containing 0.1% SDS (10mM Tris-HCl pH 7.5; 2mM EDTA; 0.1% Triton X-100; 0.1% SDS). Electrophoresis was performed by loading 30mg of total nuclear extract, and the immunoprecipitated (IP) material on NuPage 4-12% Bis-Tris gels (Novex). Protein bands were stained with Coomassie Blue.

#### Sample preparation for Mass Spectrometry (MS)

For MS analysis, IP material was digested from electrophoresis gel bands obtained from the pull-down assays. Selected gel bands were dissected and prepared using in-gel digestion protocol (Recktenwald and Hansson, 2016; van der Post et al., 2013; Shevchenko et al., 2006). Gel pieces were destained in 50% acetonitrile and reduced with 10 mM DTT at 37 °C for 30 min. Samples were alkylated with 25 mM Iodoacetoamide for 20 min at RT protected from light. The gel pieces were washed in acetonitrile between steps. Digestion was performed using 10 ng/μl trypsin (Promega) in 50 mM ammonium bicarbonate, pH 8.0 with overnight incubation at 37 °C. The peptides were extracted with a 66% acetonitrile with 0.2% formic acid solution and speed-vac to remove the organic solvents. Samples were acidified with 5% acetic acid before C18 stage-tip purification (Rappsilber et al., 2007). The peptides were resolved in 0.2% formic acid for MS analysis.

#### MS and data analysis

Analysis of MS samples was performed by LC-MS/MS using a nano HPLC system (EASY-nLC, Thermo Scientific, Odense, Denmark) coupled to a Q-Exactive HF mass spectrometer (ThermoFisher Scientific). Peptides were separated using in-house packed columns (150 x 0.0075 mm) packed with Reprosil-Pur C18-AQ 3 μm particles (Dr. Maisch, Ammerbuch, Germany). Peptide were separated with 5 to 35% gradient (A 0.1% formic acid, B 0.1% formic acid, 80% Acetonitrile) in 30 minutes. In brief, full mass spectra were acquired over a mass range of minimum 400 m/z and maximum 1600 m/z, with a resolution of at least 60,000 at 200 m/z. The 12 most intense peaks with a charge state ≥2 - 5 were fragmented with normalized collision energy of 27%, and tandem MS was acquired at a resolution of 17,500 and subsequent excluded for selection for 10 seconds. Proteins peptides were identified using MaxQuant (v1.5.7.4) in a searched against the human proteome from UniProt protein database including 75400 protein sequence entries. The modifications were set as carbamidomethylation of cysteine (fixed) and oxidation of methionine and protein N-terminal (variable).

### Bioinformatics analysis

#### RNA-seq analysis

Mapping of transcripts was done using Genome UCSC annotation set for hg38 human genome assembly. Analysis was performed using the core Bioconductor packages in R-studio v. 4.3.1. Differentially expressed genes (DEG) between the samples were identified using DESeq2 (v.1.40.2) with Benjamini-Hochberg adjustment for multiple testing.

#### ChIP-seq analysis

The fastq sequencing files were mapped to the human reference genome (hg38) using the STAR aligner (Dobin et al., 2013) with default parameters apart from setting the alignIntronMax flag to 1 for end-to-end mapping. Quality control of the sequenced material was performed by FastQC tool using MultiQC v.0.9dev0 (Babraham Institute, Cambridge, U.K.). Peak calling was performed using the HOMER findPeaks command, with 1 tag per base pair counted (-tbp 1). For peak calling in histone ChIP-seq, the option -style histone was used to find broad regions of enrichment. Peaks were filtered for the histone H3 antibody or survivin antibody IP fraction (IP) and unprocessed DNA (Input), which is a generally accepted normalization approach to identify protein-specific enrichment of DNA interaction areas (Kidder et al., 2011). A set of peaks with enrichment versus surrounding region and Input (adjusted p < 10e−5) was identified and quantified separately for each sample. Peaks were annotated with HOMER software (Heinz et al., 2010) in standard mode to the closest TSS. Peaks with overlapping localization by at least one nucleotide were merged and further on referred to as one peak. To quantify strength of binding and maintain consistency of comparison in the histone H3 samples and survivin sample, peak score was calculated by the position adjusted reads from initial peak region.

#### Tag quantification for ChIP-seq comparison

To quantify ChIP-seq tag densities from different ChIP-seq experiments, the HOMER annotatePeaks command was used, with the following parameters: -size given -noadj -pc 1. Normalized Tag Counts were calculated separately for the histone H3 ChIP-seq peaks and survivin ChIP-seq samples and presented the number of tags found at the peak, normalized to 10 million total mapped tags (Kadota et al., 2012).

#### Identification of BvCR

The R package ChIPpeakAnno, version 3.34.1 (Zhu et al., 2010) was used to identify bivalent chromatin regions, using the input ChIP-seq peaks of survivin, and the three histone H3 modifications. The function ‘findOverlapsofPeaks’ was used, with parameters restricting the maximum gap between peak ranges to zero, indicating a minimum of one bp overlap, and connected peak ranges within multiple groups as ‘merged’. The resulting set of BvCR was separated into dominant H3-BvCR. To define dominant BvCR, tags within each H3 mark of the BvCR were summed and the H3 mark that contributed the highest percentage of tags to the BvCR was designated as the dominant H3 mark in that BvCR. Peak scores of survivin within the dominant H3K4me3-BvCR, H3K27me3-BvCR, and H3K27ac-BvCR were analysed and shown as boxplots in Figure 1. Changeable BvCR were defined as those BvCR that showed a shift in the dominant H3 modification after YM155 treatment, for example, a BvCR that was initially dominant in H3K4me3 prior to YM155 treatment shifting to dominating in H3K27me3 after YM155 treatment, analysed through maximum tag percentage before and after YM155 treatment.

Matrix values to calculate the peak score per BvCR corresponding to H3K4me3, H3K27me3, and H3K27ac and survivin ChIP-seq heatmaps were generated using computeMatrix function of deepTools2, version 3.5.1 (Ramírez et al., 2016). Bedgraph files which contained the peak score corresponding to each ChIP-seq modification was used as the input to the computeMatrix function, and the bed file of BvCR was used as input to the computeMatrix function for the regions to be plotted. Using the scale-regions mode of the computeMatrix function, all BvCR, regardless of their width, were scaled to fit within a width of 500 bases, with a 2kb window upstream and downstream of the BvCR. Missing peak scores were converted to zero and a 50bp length was used for defining the score over the length of the BvCR, suggested as the default in computeMatrix function. The heatmap displays the maximum of the peak score over the length of the BvCR. The heatmaps were generated using the plotHeatmap function of deepTools2. BvCR were sorted in descending order of peak score, and the heatmap intensity was set to 50 for all the heatmaps to enable easy comparison across all histone H3 modifications. Parameters were set to default values. To examine if YM155 treatment had differential effects within and outside the BvCR, bigWigCompare function was used, which compares two bigwig files based on the number of mapped reads, where the genome is divided into several bins, and the mapped reads is counted for each bin in each of the bigwig files. Fold change was calculated for all ChIP-seq peaks of the H3 modifications or only those ChIP-seq peaks within the BvCR. If the fold change was less than 1, negative of the reciprocal of the ratio was used and interpreted as the negative fold change, as suggested by developers of the tool (Ramírez et al., 2016). The resulting bigwig file was used as the input score file for computeMatrix function and all H3 ChIP-seq peaks or the H3 ChIP-seq peaks only within BvCR were plotted using the mean of the fold change. Reference-point mode of deepTools2 was used and set to the center of the BvCR, and the fold change profile was plotted using plotHeatmap.

#### Motif enrichment analysis

Input bed files of survivin outside BvCR, and BvCR with and without survivin were used to retrieve FASTA sequences using the web service version of Regulatory Sequence Analysis Tools (Santana-Garcia et al., 2022) and the parameters of GRCh38 as the genome organism and repeats set to masked. Using these FASTA sequences, the MEME tool of MEME suite version 5.5.5 (Bailey et al., 2015) was used to identify motif sequence enrichment. Classic mode of enrichment was used, where motifs are discovered in comparison to a random model using the frequencies of the letters in the input sequence. A minimum of 10 motifs were searched, with a width between 6 and 50bp, and enrichment was performed against the known motif database of HOmo sapiens COmprehensive MOdel COllection (HOCOMOCO) version 11 FULL (Kulakovskiy et al., 2018), which contains 769 human TF binding motifs between 7bp and 25bp in width. E-value, which is an estimate of the expected number of motifs with the given log likelihood ratio (or higher), and with the same width and site count, compared to random sequences of a similar width, was used to further filter the motifs. E-value less than -100 was used, which resulted in enriched motifs for survivin and S+BvCR regions, but not for BvCR not containing survivin. The TFs found as enriched were combined and a non-redundant list was used for further analysis. FIMO tool of the MEME-Suite software, which scans sequences for motifs provided as input, was used to identify the percentage of survivin and S+BvCR regions that contained the enriched motifs. TomTom tool of MEME-Suite was used for comparison and alignment of the motifs enriched in survivin peak-containing and S+BvCR regions.

#### Peak colocalization with transcriptional regulators

To identify transcription regulators near survivin-ChIP peaks, we used the ReMap2020 database (http://remap.univ-amu.fr/) for colocalization analysis of aggregated cell- and tissue-agnostic human ChIP-seq datasets of 1034 transcriptional regulators. ReMapEnrich R-script (https://github.com/remap-cisreg/ReMapEnrich) was used for colocalization enrichment analysis.

The 4th release of ReMap (Hammal et al., 2022) present the analysis of a total of 8103 quality-controlled ChIP-seq (n=7895) and ChIP-exo (n=208) datasets from public sources (GEO, ArrayExpress, ENCODE). The hg38 human genome assembly was used for all comparisons. Two-tailed p values were estimated and normalized with the Benjamini-Yekutielli test, using the maximal allowed value of shuffled genomic regions for each dataset (n = 15), kept on the same chromosome (shuffling genomic regions parameter byChrom=TRUE). The default fraction of minimal overlap for input and catalogue intervals was set to 10%.

#### Analysis of candidate partner TFs

To identify enrichment of the BAF complex proteins within ReMap2022, we performed enrichment of our BvCR against the catalogue of all ReMap2022 ChIP-seq datasets. The BAF/SWI complex proteins, based on the list provided in a recent review (Harrod et al., 2020), were identified in this analysis and further filtered with a minimum number of overlaps > 5 and q-value of less than 0.05. Boxplots of q-significance, defined as the negative log10 of the q-value, were plotted as shown in Figure 2. To identify the overlaps of the SWI complex with BvCR, we downloaded the entire available list of ChIP-seq datasets for the SWI complex proteins and performed overlap analysis using ChIPpeakAnno with the same parameters as mentioned previously.

#### Genomic regulatory element colocalization and overlap with TF target genes

The well-curated and robust list of experimentally confirmed candidate regulatory elements spanning the human genome was obtained through the GeneHancer database version 5.9 by request (Fishilevich et al., 2017). The BED files were combined with BED files of gene bodies and 2kb upstream promoters of hg38. Genomic locations of the regulatory elements were overlapped with genome locations of overlapping BvCR using ChIPPeakAnno package with parameters mentioned previously. The entire connected gene list from GeneHancer was annotated to the overlapping dominant BvCR, and regulatory elements were retrieved. Genes from this list were selected on the dual criteria of expression (base mean >1, protein-coding) in CD4+ T cells, and transcription changes after IFNγ or YM155 treatment (nominal p-value<0.05).

#### Construction of linear model

To investigate the relationship between survivin-dependent tag deposition of histone H3 marks and survivin-sensitive gene transcription, we constructed a linear model between the minimal, median, and maximal values of the observed tag deposition change after YM155 treatment, and the observed transcriptional change through RNA-seq. A line of best-fit, using the min-max approach, was drawn using the coordinates of these three points. Using the slope and the intercept of this line, we predicted the tag percentage for each observed fold change and calculated the difference between the predicted and observed fold changes. If this difference fell within one standard deviation of all the predicted vs observed fold changes, these genes were included in the model. For these included genes, Spearman correlation was calculated using ‘stat_cor’ function in the ‘ggpubr’ R package. Correlation modelling was performed for either all BvCR or BvCR containing survivin colocalization i.e., S+BvCR. Correlations were calculated separately for upregulated genes and downregulated genes. Radar plots were generating using the ‘ggradar’ R package.

#### Pathway enrichment analysis

Pathway enrichment analysis was performed using ShinyGO version 0.77 (Ge et al., 2020), restricting the search space to the pathway size of 100 to 1500 genes in human GO: Biological Process (GO: BP) terms, and FDR < 0.05. The top 20 enriched terms were selected and further filtered based on the redundancy in term definition and the number of genes annotated to the enriched terms. For pathway enrichment analysis in BRG1^hi^ and BRG1^lo^ cells through the WGCNA approach, the web version of Enrichr (Chen et al., 2013) was used, restricting the search space within GO: Biological Process (GO: BP) and Reactome 2022 (reactome.org).

#### DNA Damage Response network analysis

The network map detailing subprocesses of the DNA damage response (DDR) was retrieved from a recently published affinity purification-mass spectrometry study that catalogued the protein-protein interactions of 21 central DNA damage response proteins (Kratz et al., 2023). This network consists of 605 genes which cover 109 hierarchical categories in total. Of this list, genes controlled by the BvCR were extracted. The dominant H3 mark in each category was estimated by the average tag percentage of the H3 mark within the BvCR.

#### Weighted Gene Correlation Network Analysis (WGCNA)

To identify modules of genes that associated with BRG1-high and BRG1-low cells, we performed the Weighted Gene Correlation Network Analysis using the R package WGCNA version 1.72-5 (Langfelder and Horvath, 2008). The matrix of normalized gene expression values of genes connected to BvCR and enriched in at least one of the pathways was used as input for WGCNA. A soft-threshold power of 10 was chosen as it was the lowest power at which the fit index reached 90%. The ‘signed’ network type was used to identify genes positively correlated (Pearson correlation) in samples having high and low BRG1 expression. A minimum module size of 30 was used with ‘mergeCutHeight’ set to 0.25. This resulted in three modules of which one module contained only 4 genes. We explored the module-gene relationships and found that this module showed correlation profiles similar to that of the module containing positively correlated genes in BRG1-low samples. Hence, we merged the genes within these modules and performed pathway enrichment analysis of the resulting two modules as detailed above.

#### Compositional analysis of cBAF and pBAF subunits

The peptides from the survivin binding custom designed microarray (n>5250, PEPperCHIP, Heidelberg, Germany) comprised a scikit-learn python library to implement the machine learning process, in which the peptides were divided into equally large training and test sets (Jensen et al., 2023). Based on the functional composition of the protein, defined by the presence of atomic groups (C, CH, CH_2_, CH_3_, hydroxyl, phenyl, carboxyl, amide, sulfhydryl, etc.) rather than the sequence of amino acids, we develop a strategy for predicting fitness of a given protein/peptide to survivin in biological and a chemical context (Jensen et al., 2023; Anindya et al., 2024) with the following adjustments. The multilayer perceptron classifier comprised two intermediate layer neurons, and C-Pos encoding was employed to interpret 15 amino acid-long peptides. Peptides with a C-terminal cysteine were excluded from the training data, and 90% of the original survivin peptide microarray data (Anindya et al., 2022) was utilized for training purposes.

The sequence data of both cBAF and pBAF subunits was segmented into 15 amino acid-long peptides with 10 amino acid overlaps. Subsequently, the trained model was utilized to predict the wild-type sequence, enabling the calculation of R_bind_ (the ratio of predicted binding peptides to the total number of generated peptides). M_bind_(n) values were computed for each amino acid position by mutating the position to all other amino acids and assessing whether the machine learning model predicted survivin binding. M_bind_(n) was defined as the ratio of the total number of predicted mutant peptides (for a specific position) to the total generated mutations.

#### Data analysis and visualization

Statistical analysis was performed using R-studio (version 4.3.1). Heatmaps were visualized using the ComplexHeatmap R package version 2.16.0 (Gu et al., 2016). Schematic visualizations were created using biorender.com.

## Data availability statement

RNA and ChIP sequencing data (raw data and processed files) of CD4+ T cells that support the findings of this study have been deposited in NCBI GEO. Survivin ChIP-seq: GSE190354; H3K27me3 ChIP-seq ± YM155 treatment: GSE216818; H3K27ac ChIP-seq ± YM155 treatment, GSE216817; H3K4me3 ChIP-seq ± YM155 treatment, GSE216820. RNA-seq: 1) CD4^+^ T cells RA patients, GSE190349; 2) human CD4^+^ T cells treated with IFN, IFN+YM155, and control, GSE190351; 3) CD4^+^ T cells of RA patients treated with JAK-inhibitors and untreated, GSE201669.

The mass spectrometry proteomics data of survivin affinity immunoprecipitated material have been deposited to the ProteomeXchange Consortium via the PRIDE partner repository (Perez-Riverol et al., 2022) with the dataset identifier PXD049683.

Other RNA-seq data sets: CTLA4 fusion protein abatacept treatment GSE121827, IL6 receptor blocking antibody tocilizumab treatment GSE113156, and methotrexate treatment GSE176440. CD4+ T cell clusters in RA synovial tissue by CITE-seq, https://doi.org/10.1038/s41586-023-06708-y

All other relevant data supporting the key findings of this study are available within the article and its Supplementary Information files from the corresponding author upon reasonable request. Source data are provided in this paper.

## Acknowledgements

We would like to thank the research nurses Anneli Lund and Marie-Louise Andersson at the Rheumatology Clinic, Sahlgrenska University Hospital, Gothenburg, for their help with blood sampling. We would like to thank Nina Oparina for introducing us to ChIP-seq analysis. We also thank all patients with RA, who participated in this study. This work has been funded by grants from the Swedish Research Council (MB, 2017-03025 and 2017-00359), the Röntgen-Ångström Cluster Framework of the Swedish Research Council (GK, 2015-06099), the Swedish Association against Rheumatism (MB, R-566961, R-751351 and R-860371), the King Gustaf V’s 80-year Foundation (MB, FAI-2018-0519 and FAI-2020-0653), the Regional agreement on medical training and clinical research between the Western Götaland county council and the University of Gothenburg, Sweden (MB, ALFGBG-717681, ALFGBG-965623). The authors declare that the funding sources have no role in study design; in the collection, analysis, and interpretation of data; in the writing of the report; and in the decision to submit the article for publication.

## Author contributions

V. Chandrasekaran, M.I. Bokarewa, G. Katona, conceived the study. K. Andersson, M. Erlandsson collected materials, and performed laboratory work. M.J. Garcia-Bonete performed mass spectrometry experiments and proteomic data analysis. V, Chandrasekaran, M. Erlandsson, K. Andersson, and E. Malmhäll-Bah performed bioinformatic analysis. S. Li performed molecular docking analyses. G. Katona performed machine learning prediction modelling. P. Johansson provided reagents and input in drafting of manuscript. V. Chandrasekaran, M.I. Bokarewa drafted the article. All authors discussed and helped interpret the data and provided feedback during the preparation of the article.

