## Supplementary figures for "SURVIVIN IN SYNERGY WITH BAF/SWI COMPLEX BINDS BIVALENT CHROMATIN REGIONS AND ACTIVATES DNA DAMAGE RESPONSE IN CD4+ T CELLS"

### Supplementary Figure S1

S1A. Box plots of histone H3 tag deposition within the bivalent chromatin regions (BvCR) dominant by H3K4me3, H3K27me3 and H3K27ac

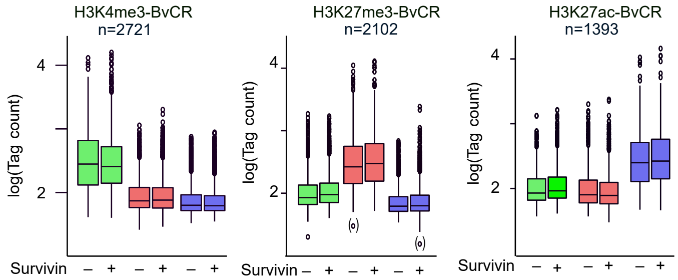


S1B. Box plots of histone peak scores within BvCR dominant by H3K4me3, H3K27me3 and H3K27ac. Kolmogorov-Smirnov test p-values are shown.

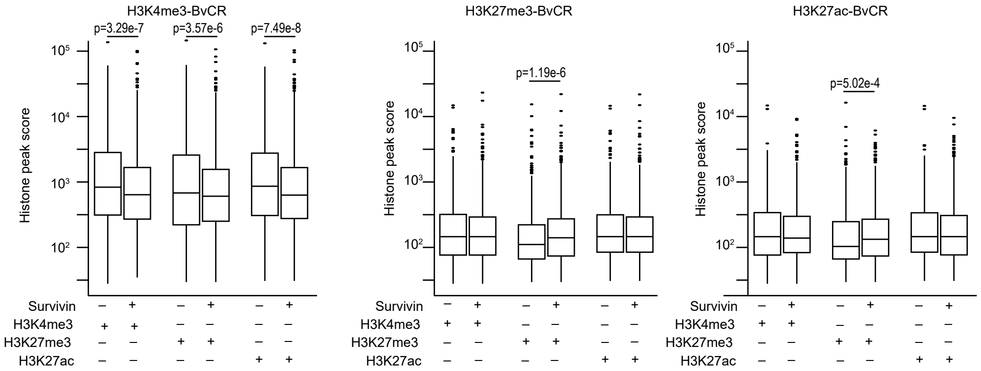


S1C. Box plot of percentage tag change for histone peaks within H3K27me3- and H3K27ac-BvCR, after YM155 treatment. Mann-Whitney test p-values are indicated.

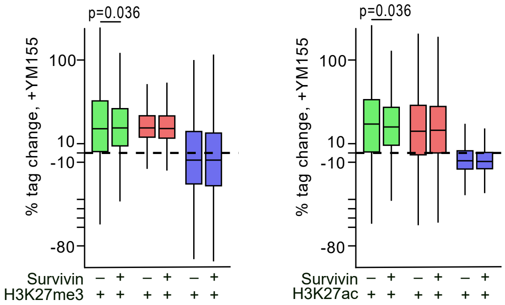

### Supplementary Figure S2

23 Feb 2024

S2A. Forest plot of probability of BvCR to be changeable (Ch) in YM155-treated CD4 cells and genes connected to BvCR to be differentially expressed (DEG) in CD4+ cells treated with IFNγ or IFNγ+YM155.

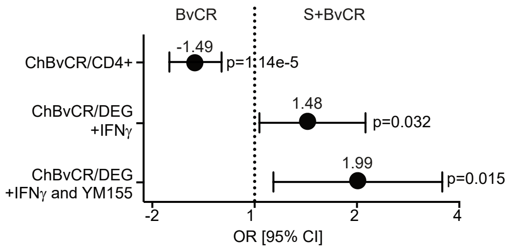


S2B. Scatter plot of correlation between change in deposition of H3K4me3 and H3K27me3 tags and change in transcription of DEG after treatment with IFNγ or IFNγ+YM155 in survivin-positive H3K4me3-BvCR. Spearman ρ are indicated.

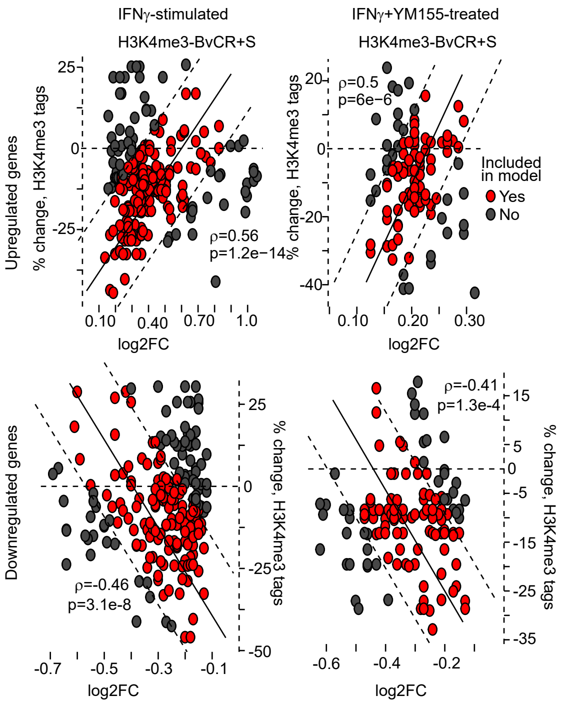


S2C. Radar plot of Spearman's ρ correlations between tag change in H3K4me3 and H3K27me3 deposition in H3K27me3-BvCR and transcription change of DEG in CD4⁺ cells treated with IFNγ or IFNγ+YM155. Arrows indicate direction of transcription change

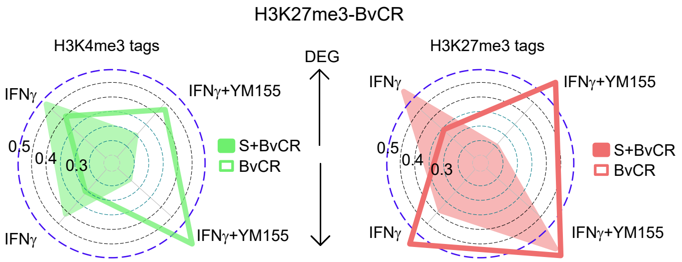

**Supplementary Figure S3** 23 Feb 2024

S3A. Bar plot of frequency of genes connected to BvCR in the enriched pathways. To the right is the Venn diagram of IFNγ- and survivin-sensitive genes connected to H3K4me3-BvCR and annotated to the DNA damage response pathway (GO:0006974)

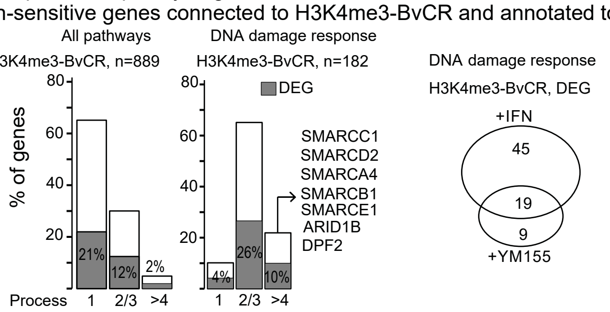


S3B. Heatmap of normalized tag deposition in BvCR connected to DEG treated with IFNγ+YM155. Filled squares indicate colocalization of survivin (S) in the BvCR. Genes connected to multiple BvCR are marked in bold

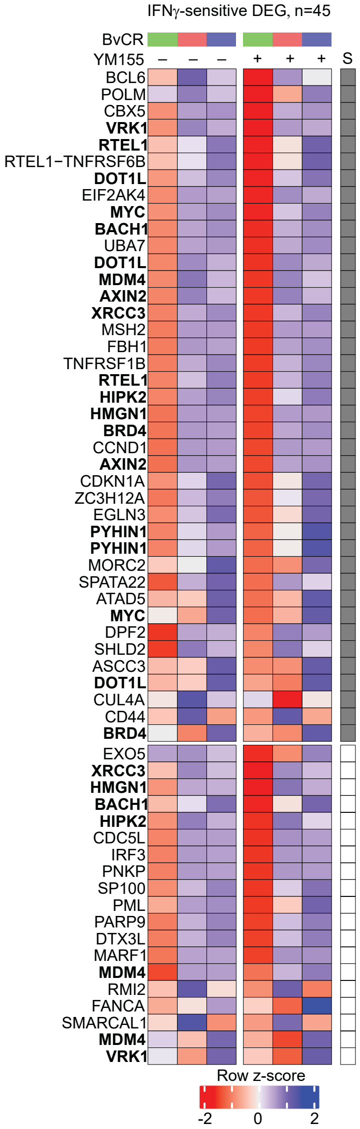


S3C. Heatmap of transcription change of DEG annotated to DNA Damage Response (DDR) pathway. Asterisks indicate RNAseq nominal p-values - * < 0.05, ** < 0.01, *** <0.001.

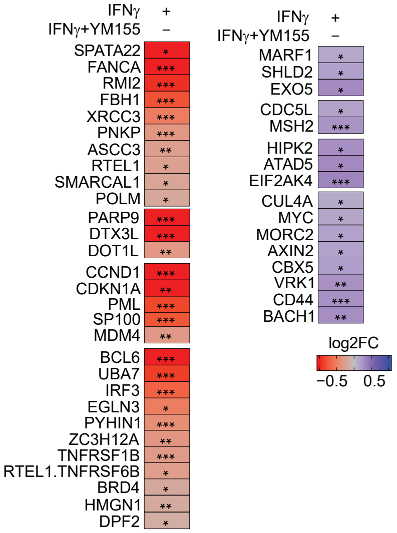


S3D. Box plot of quantified tag deposition in survivin-sensitive and IFNg-sensitive genes annotated to DNA damage response and connected to H3K4me3-BvCR. Mann-Whitney p-values are indicated

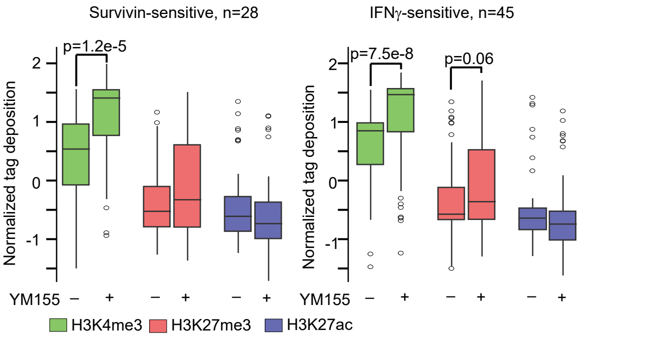

**Supplementary Figure S4** 23 Feb 2024

S4. Genomic maps of the DNA repair gene loci MSH6, FANCI, SMC3, PIAS4, and MRE11. Filled black and red boxes indicate cis-RE connected to the gene, as determined by GeneHancer. Distance to TSSs is shown. Black filled peaks underneath cis-RE indicate the positions of survivin-ChIP and histone H3-ChIP peaks. Colored peaks indicate the change in tag deposition for H3K4me3 (green) and H3K27me3 (red) after YM155 treatment, scaled to enable direct comparison between the two modifications.

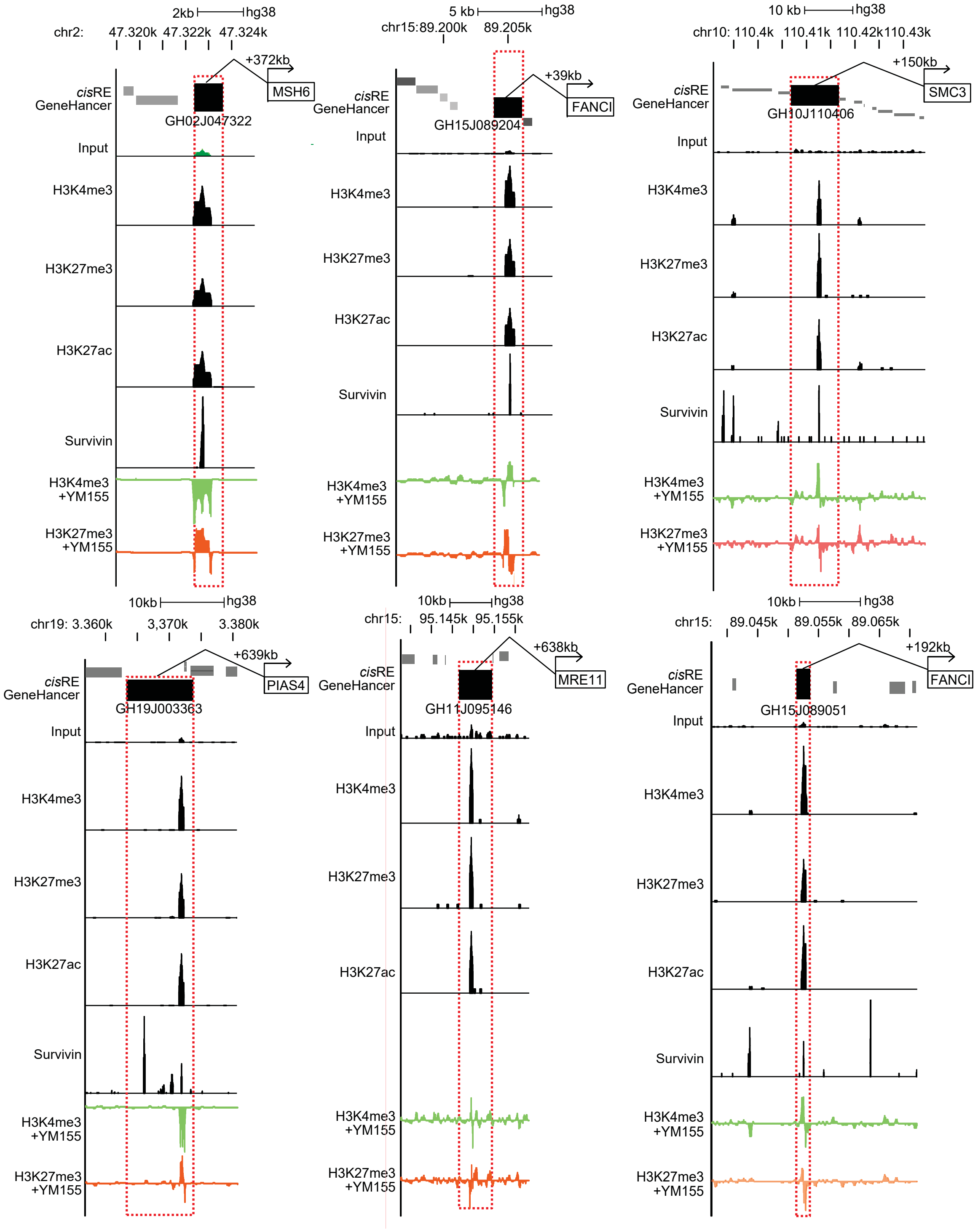

### Supplementary Figure S5

23 Feb 2024

S5A. Frequency of overlapping BvCR with cBAF and PBAF complex subunits retrieved from ReMap2022 database. Fisher test p-values are indicated.

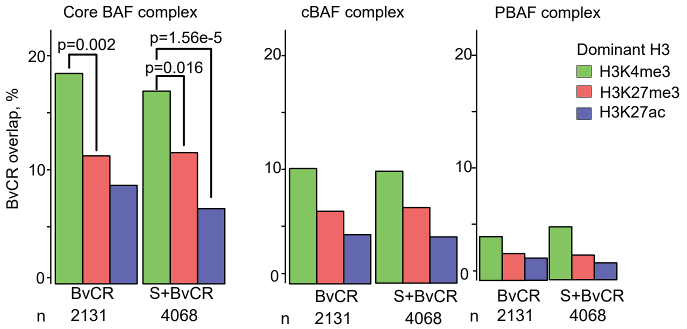


S5B. Distribution of survivin binding probability across the protein sequence of SMARCC2, SMARCD1, and SMARCE1. $M_{bind}(n)$ value indicate the fraction of mutations compatible with a survivin binding to the residue, defined by the functional composition of atomic group (Anindya et al, 2024). “1” indicates a region predicted to bind survivin even if the position is mutated to any other amino acid. “0” indicates no mutation can convert the site to survivin binding region. Uniprot IDs of the proteins are indicated in brackets.

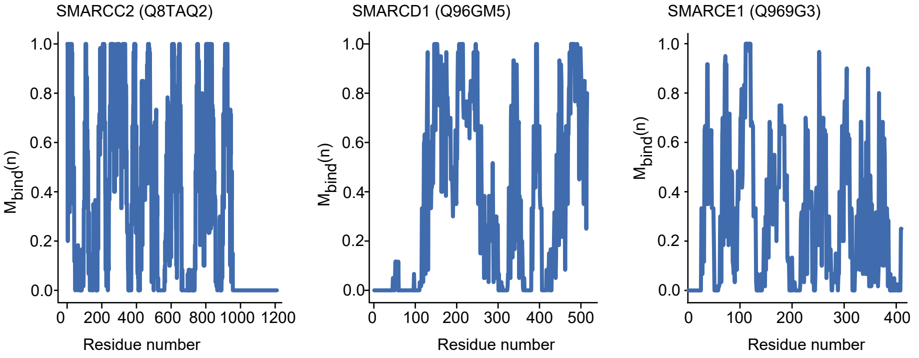


S5C. The ribbon diagram of interaction between survivin and the PBAF complex predicted by docking modelling. Residues of the PBAF complex involved in the interaction with survivin are depicted in the surface representation.

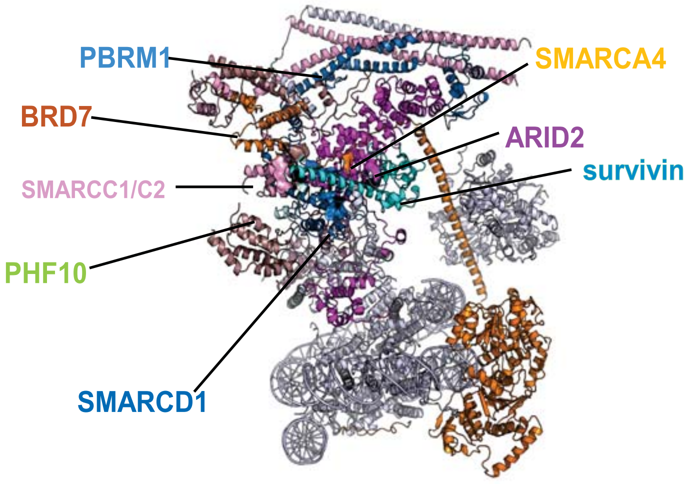

### Supplementary Figure S5



S5D. Table of amino acid residues within cBAF and PBAF subunits that bind survivin. The residue numbering conventions introduced in the PDB entries 6ltj (He et al, 2020) and 7vdv (Yuan et al, 2022) were utilized. Amino acid residues were identified by the docking analysis. $M_{bind}(n)$ values are predicted by compositional analysis of each residue.

| Subunits | Conventional BAF | | Subunits | Polybromo BAF | |
|---|---|---|---|---|---|
| | Residues | $M_{bind}(n)$ | | Residues | $M_{bind}(n)$ |
| SMARCA4 | **380-HIS**<br>381-ARG<br>384-GLU<br>396-LEU<br>399-LYS<br>403-GLU | **0.7**<br>0.2<br>0.2<br>0.1<br>0.1<br>0.2 | SMARCA4 | **437-LYS**<br>**438-ALA** | **0.5**<br>**0.7** |
| SMARCC1/C2 | 600-GLU<br>**643-PRO**<br>**646-ASP**<br>**647-PRO**<br>**650-GLU**<br>**651-ASP**<br>**656-LEU**<br>682-SER<br>683-VAL<br>700-PHE<br>701-SER<br>702-LYS<br>703-MET | 0.2<br>**1.0**<br>**1.0**<br>**1.0**<br>**1.0**<br>**0.7**<br>**0.8**<br>0.0<br>0.0<br>0.0<br>0.1<br>0.1<br>0.1 | SMARCC1/C2 | **614-GLU**<br>**615-MET**<br>**616-TYR**<br>**617-LYS**<br>**618-ASP** | **1.0**<br>**1.0**<br>**0.7**<br>**1.0**<br>**0.8** |
| SMARCD1 | **480-GLU**<br>**483-PHE**<br>**485-PRO** | **1.0**<br>**1.0**<br>**1.0** | SMARCD1 | **505-GLU**<br>**508-GLN**<br>**509-ALA**<br>**510-LEU** | **0.6**<br>**0.8**<br>**0.9**<br>**0.5** |
| DPF2 | 65-GLY<br>66-LEU<br>67-ALA<br>68-SER | 0.1<br>0.3<br>0.4<br>0.4 | PHF10 | **280-ASN**<br>**281-THR** | **0.8**<br>**0.6** |
| ARID1A | **1722-ARG**<br>**1726-GLU**<br>**1732-LYS**<br>**1738-ASP**<br>**1833-ARG**<br>1853-GLU<br>1855-ILE<br>1862-LYS | **1.0**<br>**1.0**<br>**1.0**<br>**1.0**<br>**0.9**<br>0.1<br>0.2<br>0.2 | ARID2 | 187-SER<br>188-LYS | 0.0<br>0.0 |
| | | | BRD7 | 378-LEU<br>379-GLN<br>380-SER | 0.0<br>0.0<br>0.0 |
| | | | PBRM1 | 1599-PRO<br>1601-THR<br>1604-LEU | 0.0<br>0.1<br>0.1 |

**Supplementary Figure S6** 23 Feb 2024

S6A. The table of clinical characteristics of healthy controls and patient material used in this study.

| | Healthy Controls | GSE190349 | GSE201669 | GSE176440 | GSE121827 | GSE113156 |
|---|---|---|---|---|---|---|
| n | 40 | 24 | 33 | 28 | 14 | 6 |
| Treatment | N/A | N/A | JAKi | Methotrexate | Abatacept | Tocilizumab |
| Mean age, years (range) | 51.6 (26-77) | 53.8 (25-73) | 54.3 (23-73) | 59.5 (47.5-67) | 37.7 (24-58) | 65 (52-67) |
| Female gender, % | 97.5 | 100 | 100 | 71.4 | 78.6 | 66.7 |
| Disease duration, median (IQR), months | N/A | 66 (20-134) | 126 (72-228) | 84 (36-288) | N/A | 67 (14-293) |
| ESR, median (IQR), mm/h | N/A | 19.5 (8.7-25.2) | 11 (5-20) | 31 (14-46) | N/A | 41 (13-62) |
| ACPA-positive, n (%) | N/A | 12 (50%) | 18 (54%) | 26 (92.9) | N/A | 4 (66.7) |
| RF-positive, n (%) | N/A | 12 (50%) | 19 (57%) | 25 (89.2) | N/A | 4 (66.7) |

S6B. DNA Damage Response (DDR) network map of upregulated (red) and downregulated (blue) DEG in BRG1$^{hi}$ cells in RA patients.

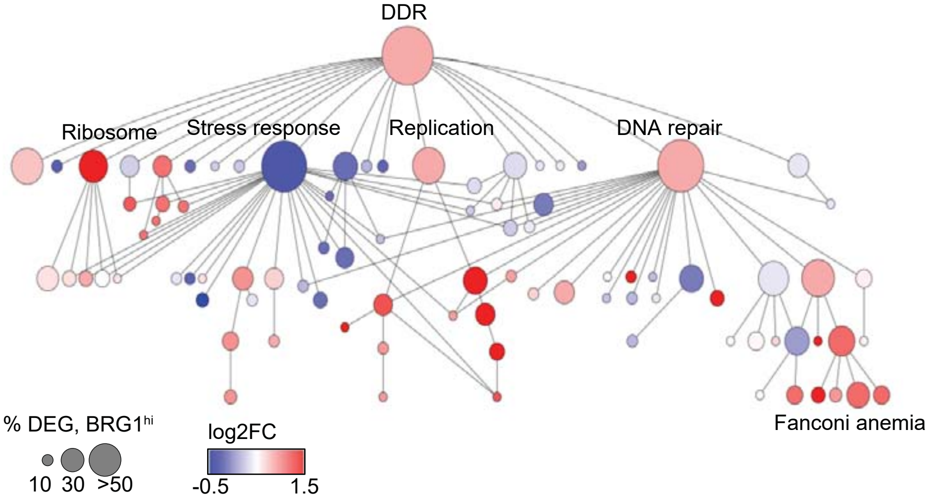


S6C. The table of primers used for qPCR analysis in this study.

| | Forward | Resverse |
|---|---|---|
| BIRC5 | GACCACCGCATCTCTACATTC | TGCTTTTTATGTTCCTCTATGGG |
| FANCI | CCTCCAAGGGAAGCAGAAAG | TCTCCTTGCTGTCCTACCTT |
| MRE11 | ACAAGGAGGAGAAAGATGCCA | TCATAGCCTCACGGACTTCAT |
| MSH6 | AGCCCTCAGAGCCAGAAGA | AATTCCACATCAGAGCCACC |
| PFKFB3 | CCTACAACTTCTTCCGCCCC | CCGCAATTTGTCCCCCTTCT |
| SMARCA4 | ACCAGAAGCAGAGCCGCAT | TCAATGGTCGCTTTGGTTCG |
| SMARCC1 | AGCTGTTTATCGACGGAAGGA | CTGAAGAAGCTGCACCACC |
| SMARCE1 | ACCATCTTATGCCCCACCTC | TCCCAGCCTGTAGTTGTTGT |
